# Cannabinoid Receptor Signaling is Dependent on Sub-Cellular Location

**DOI:** 10.1101/2024.03.21.586146

**Authors:** Alix Thomas, Braden T Lobingier, Carsten Schultz, Aurélien Laguerre

## Abstract

G protein-coupled receptors (GPCRs) are membrane bound signaling molecules that regulate many aspects of human physiology. Recent advances have demonstrated that GPCR signaling can occur both at the cell surface and internal cellular membranes. Our findings suggest that cannabinoid receptor 1 (CB1) signaling is highly dependent on its subcellular location. We find that intracellular CB1 receptors predominantly couple to Gαi while plasma membrane receptors couple to Gαs. Here we show subcellular location of CB1, and its signaling, is contingent on the choice of promoters and receptor tags. Heterologous expression with a strong promoter or N-terminal tag resulted in CB1 predominantly localizing to the plasma membrane and signaling through Gαs. Conversely, CB1 driven by low expressing promoters and lacking N-terminal genetic tags largely localized to internal membranes and signals via Gαi. Lastly, we demonstrate that genetically encodable non-canonical amino acids (ncAA) offer a solution to the problem of non-native N-terminal tags disrupting CB1 signaling. We identified sites in CB1R and CB2R which can be tagged with fluorophores without disrupting CB signaling or trafficking using (*trans*-cyclooctene attached to lysine (TCO*A)) and copper-free click chemistry to attach fluorophores in live cells. Together, our data demonstrate the origin of location bias in cannabinoid signaling which can be experimentally controlled and tracked in living cells through promoters and novel CBR tagging strategies.

## Introduction

G-protein coupled receptors (GPCRs) are essential for regulating human development and physiology, and their perturbation can have dramatic effects on onset and disease progression. As a result, 30% of FDA approved drugs target GPCRs(1). Although there are over 200 structurally distinct GPCRs, they all signal via a few G-proteins. Recently, our understanding of how signal specificity is achieved by GPCRs and how it can be translated to therapeutic intervention has greatly increased(2, 3). Among other significant advancements, the concept of functional selectivity acknowledges that the same receptor can produce different cellular outcomes by modulating the specificity and timing of downstream events(4). This functional selectivity is thought to occur either through biased agonism (i.e, the ability of a GPCR to adopt different receptor and scaffold conformation based on its ligand)(5, 6), or through location bias (i.e, the ability to signal from different subcellular localizations)(7–9). Emerging data are giving rise to a new signaling model where ligands bind and activate GPCR both at the cell surface and at internal membranes. For example, the B1 adrenergic receptor can stimulate an intracellular Gαs-mediated cAMP signal from the Golgi apparatus, thereby significantly contributing to the increase in cAMP levels (10), while the opioid receptors mu and delta can also couple to Gαi/o at the Golgi apparatus(11). This location bias can dramatically modulate the activity of therapeutics as it requires the drug to either be actively transported or passively diffusing to the receptor’s sub-localization.

Cannabinoid receptor 1, a rhodopsin-like G protein coupled receptor, is generally described as a Gαi/o coupled receptor although its specificity for Gαi/o was challenged years ago with evidence that the receptor is able to couple to numerous G proteins in different cell types(12, 13). It was shown by Diez-Alarcia *et al* that the cannabinoids THC, WIN55 and ACEA can stimulate not only the Gαi/o pathway but also Gαs, Gαq and Gα12/13 in a ligand dependent fashion(14). These findings align with reports that stimulation of CB1R in different areas of the brain or in peripheral tissues yields diverse outcomes(15). In addition to the diversity of CB1R coupling to G proteins, there is also heterogeneity in its expression and subcellular location. The expression level of CB1R is very high in the brain(16), but varies widely across other cell types (17, 18). This fluctuating expression level was shown to have implication in the receptor ligand binding and G protein activation(19) yet the reasons for these signaling differences remain unclear. CB1R has been described to reside and signal from intracellular compartment(15) such as late endo-lysosomes(20) and mitochondria. Here, it was shown to modulate mitochondrial respiration, intra-mitochondrial cAMP levels and PKA activity(21). To date it remains unclear if this differential coupling of the CB1R to G proteins originates from cell type-specific fluctuations in its expression level or represents a general principle in which signaling is influenced by location bias. Given the recent appreciation for location bias in GPCR signaling, we hypothesized that the expression level and sub-cellular localization of the CB1R plays a critical role in determining its affinity and interaction profile of G-proteins.

In order to evaluate the impact of both expression level and location bias of CB1R on downstream signaling, we leveraged approaches commonly employed in GPCR molecular pharmacology. We used higher (CMV) and lower (UBC) expressing promoters to heterologously drive CB1R expression in HeLa cells and we also examined the effects of the commonly utilized N-terminal signal sequence FLAG (SSF) tag for monitoring GPCR expression and trafficking (22). We selected HeLa cells, which lack endogenous CB1R yet possess the necessary downstream effectors for CB1 signaling, as a commonly used cellular model. We found that lower expression levels predominantly localized CB1R within internal organelles and signaled via Gαi, contrasting with the prevalent plasma membrane localization and Gαs coupling at higher expression levels. Furthermore, utilizing the SSF tag in conjunction with low CB1R expression mimicked the receptor’s predominant plasma membrane localization previously observed at higher expression levels. This model suggests that the plasma membrane pool of CB1R stimulates cAMP production via Gαs coupling, whereas the endo-membrane pool reduces cAMP levels through Gαi/o coupling. To visualize CB1R without interfering with its localization, we incorporated a single non-canonical amino acids in CB1R (*trans*-cyclooctene lysine (TCO*A)) and attached fluorophores by ultrafast copper-free click chemistry in live cells(23).

## Results

### Expression level and N-terminal tagging direct the sub-cellular localization of CB1R

Immunofluorescence analysis revealed the absence of endogenous cannabinoid receptor 1 (CB1R) expression within our HeLa cell line (Figure S1.2A). We established two distinct expression systems: _CMV_CB1 and _UbC_CB1, driven by the stronger CMV and weaker UbC promoters, respectively(24). Confocal microscopy imaging delineated disparate subcellular localization patterns between the two expression systems. _UbC_CB1 primarily localized within endomembranes, notably the endoplasmic reticulum (ER), Golgi apparatus, as well as a minor fraction at the plasma membrane. Conversely, _CMV_CB1 predominantly localized to the plasma membrane, with a residual presence observed at the Golgi apparatus and ER (Figure 1A, S1.1A,B,C). Thus, in our heterologous expression system, subcellular localization of CB1R was largely controlled by promoter strength.

**Figure 1.**
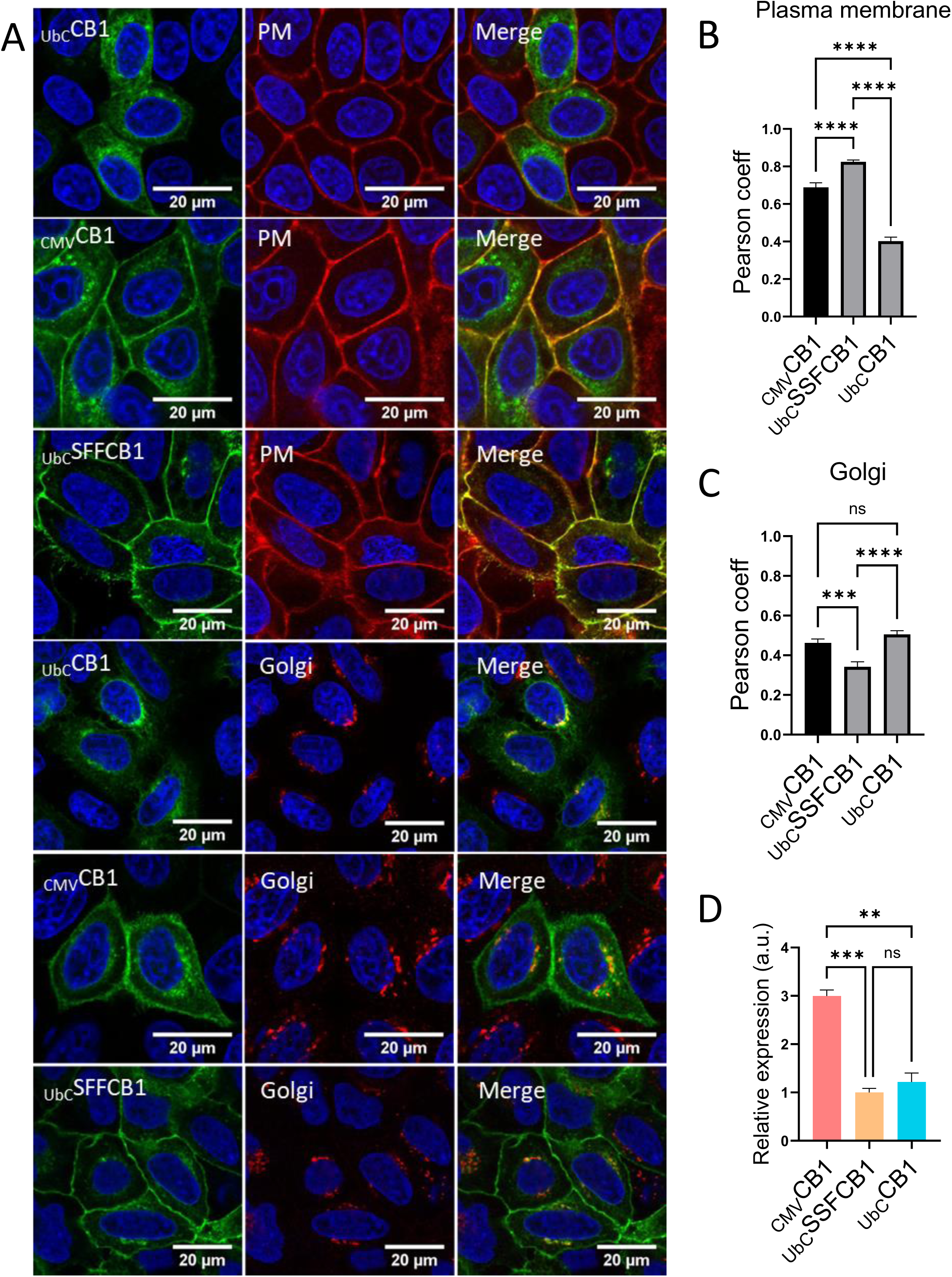
Expression level and N-terminal modification direct CB1R’s sub-cellular localization and impact downstream cAMP signaling. **A.** Confocal micrographs showing HeLa Kyoto cells transfected with _UbC_CB1R, _CMV_CB1R or _UbC_SSFCB1R. Cells were fixed, permeabilized and co-immuno-stained with a CB1R antibody (left), and respective PM and Golgi organelle marker CellBrite and GM130 antibody (middle). **B.** Colocalization measurement in HeLa cells using Pearson coefficient of immunolabeled _UbC_CB1R, _CMV_CB1R or _UbC_SSFCB1R with the plasma membrane stain CellBright. **C.** Colocalization measurement in HeLa cells using Pearson coefficient of immunolabeled _UbC_CB1R, _CMV_CB1R or _UbC_SSFCB1R with the immunolabeled Golgi apparatus. **D.** Bar graphs showing n=3 replicates of expression levels of _UbC_SSFCB1R-APEX, _UbC_CB1R-APEX and _CMV_CB1R-APEX. _CMV_CB1-APEX / SSFCB1-APEX p < 0.005. _CMV_CB1-APEX / _UbC_CB1-APEX p < 0.01. SSFCB1R-APEX / _UbC_CB1R-APEX was non-significantly different.

We next wanted to employ tagging methods to monitor CB1R expression and agonist-induced trafficking. We tagged CB1R at its N-terminus with the signal sequence FLAG (SSF) tag as antibody epitope tags like SSF are a common approach in the molecular pharmacology field to monitor GPCR expression, trafficking, and improve delivery to the cell surface(22, 25, 26). We introduced a modified construct called _UbC_SSF-CB1, where a signal sequence flag (SSF) tag was added to the N-terminus of CB1R. We found that the SSF tag caused _UbC_SSF-CB1 to largely be localized to the plasma membrane and mostly excluded from endo-membranes (Figure 1A,B,C, S1.1A,D). To ensure that the SSF tag’s effect on localization was not due to higher expression levels relative to _UbC_CB1, we used a CB1-APEX2 fusion protein to quantify whole cell receptor expression(27). We found that the expression level of _UbC_SSF-CB1 was comparable to that of _UbC_CB1, while _CMV_CB1 showed a three-fold increase in expression compared to when the SSF tag was present (Figure 1D, S1.2B). It is unclear how the SSF tag directs CB1R to the plasma membrane, the original goal of the tag was to enhance cell surface delivery of GPCRs by introducing a non-native and cleavable signal sequence from influenza hemaglutinin (22). We found that the SSF tag can be cleaved in live cells by trypsin, possibly due to the uniquely long N-terminal tail of CB1R (Figure S1.2C). This supports the hypothesis that the N-terminal tail plays a significant role in CB1R trafficking and expression regulation, as was postulated before (28). In conclusion, our results demonstrate that the localization of CB1R is highly dependent on expression level and N-terminal tagging.

### Sub-cellular localization of CB1R impacts downstream signaling via cAMP

Our investigation aimed to discern how CB1R’s subcellular localization might affect cell signaling. CB1R is well known to predominantly signal via Gαi/o proteins at the plasma membrane but has also been reported to couple to Gαs and Gαq(14, 29). These interactions lead to alterations in cAMP and calcium levels, respectively. Recent studies have suggested that the intracellular pool of CB1R may play a role in cell signaling, prompting an exploration into the potential signaling processes originating from intracellular membranes (17). To measure changes in cAMP levels upon CB1R activation, we employed a genetically encoded EPAC-based FRET sensor (30). Cells were co-transfected with different CB1R constructs (_CMV_CB1, _UbC_CB1, and _UbC_SSF-CB1) and pretreated with forskolin (50µM) before treating with the CB1R full agonist WIN55,212 (10µM). Cells expressing _UbC_CB1, which predominantly localizes CB1R to endo-membranes, exhibited a decrease in cAMP levels after WIN55,212 treatment, indicating Gαi/o coupling. In contrast, cells expressing _CMV_CB1, mainly located at the plasma membrane, showed an increase in cAMP levels, suggestive of Gαs coupling (Figure 2A,C).

**Figure 2.**
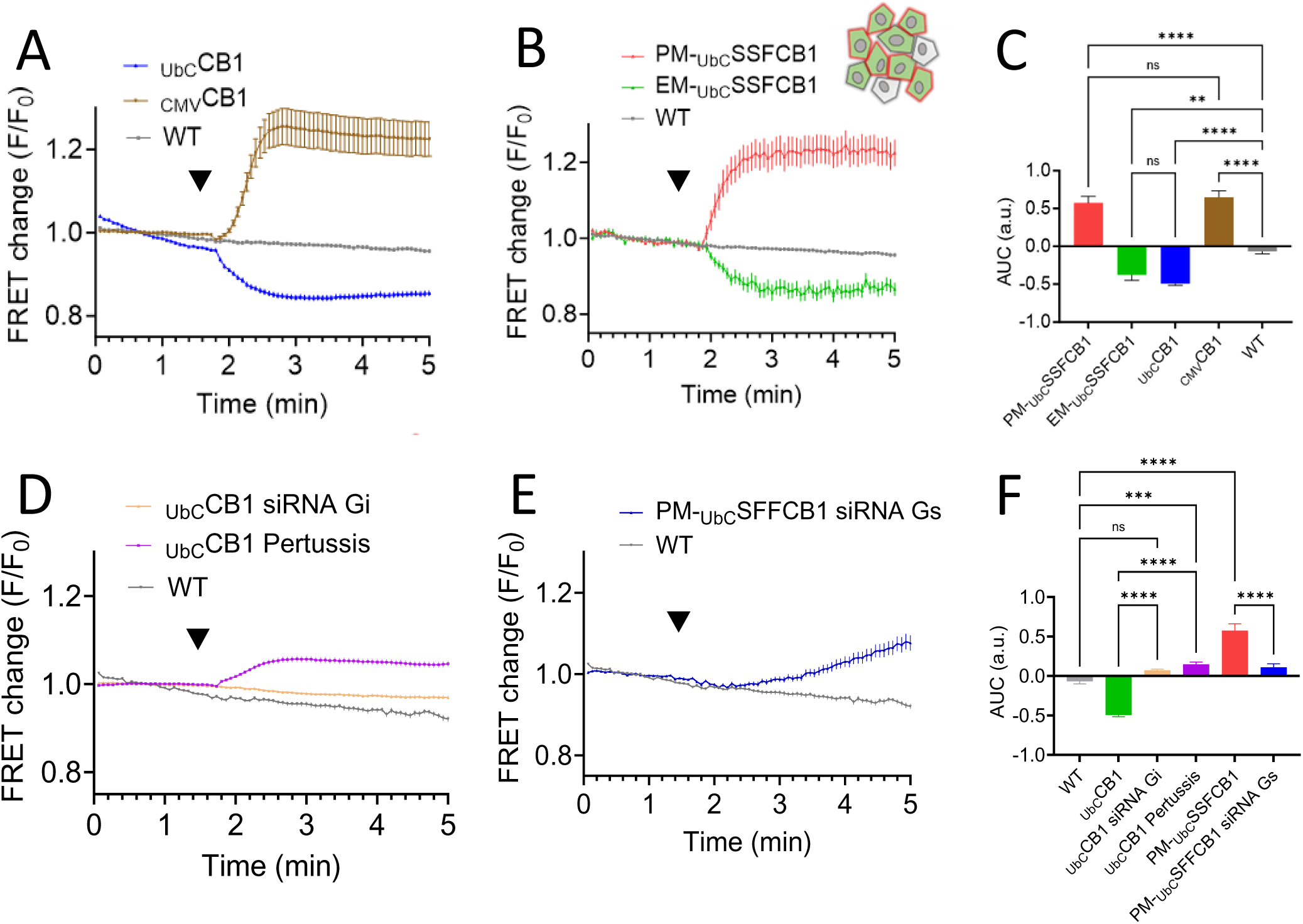
Location bias drives emCB1 to couple to Gαi/o and pmCB1 to couple to Gαs. **A.** Average of 111, 63, and 57 traces showing FRET changes of the EPAC-based FRET sensor in _UbC_CB1R-expressing, _CMV_CB1R-expressing or non-expressing (WT) HeLa cells, respectively. **B.** Average of 67, 46 and 57 traces showing FRET changes of the EPAC-based FRET sensor in SSF_UbC_CB1R-transfected HeLa cells with (PM) or without (EM) tagging with the Flag antibody or not transfected (WT), respectively, after treatment with the agonist WIN55,212-2 (10 µM). **C.** Bar graphs showing area under the curve of the EPAC-based FRET sensor after WIN55,212-2 stimulation (10uM) in PM-SSF_UbC_CB1R expressing cells, EM-_UbC_SSFCB1R, _UbC_CB1R expressing cells and WT non-expressing cells. PM-_UbC_SSFCB1R / WT p < 0.001, EM-_UbC_SSFCB1R / WT p < 0.001, _UbC_CB1R / WT p < 0.001. **D.** Average of 179, 176 and 84 traces showing FRET changes of the EPAC-based FRET sensor in _UbC_CB1-expressing HeLa, pre-treated with pertussis toxin 24h prior to experiment, pre-transfected with Gαi/o siRNA 72h prior to experiment or WT-not expressing HeLa respectively, after treatment with the agonist WIN55,212-2 (10 µM). **E.** Average of 116 and 84 traces showing FRET changes of the EPAC-based FRET sensor in SSFCB1R-transfected HeLa cells tagged (PM) with the Flag antibody, pre-transfected with Gαs siRNA 72 h prior to the experiment or WT-not expressing HeLa, respectively, after treatment with the agonist WIN55,212-2 (10 µM). Cells in all experiments were pre-treated with forskolin (50 μM). **F.** Bar graphs showing area under the curve of the EPAC-based FRET sensor after WIN55,212-2 stimulation (10 µM) in WT non-expressing cells, PM-_UbC_SSFCB1R-expressing cells pre-transfected with Gαs siRNA and, _UbC_CB1R-expressing cells pre-transfected with Gαi/o siRNA. PM-_UbC_SSFCB1R siRNA Gαs and _UbC_CB1R siRNA Gαi/o versus WT were not significant. Cells in **A** to **F** were pre-treated with forskolin (50 µM).

We next sought to take advantage of the fact that our _UbC_SSF-CB1 resulted in a mixed population in which most cells had CB1R at the plasma membrane but a subset retained CB1R internally (S2.1). We labeled _UbC_SSF-CB1 receptors with a non-cell-permeant 647Alexa-M1-FLAG antibody prior to the experiment for sub-cellular localization analysis. Using the flag tag to identify plasma membrane CB1R and immunostaining post cell-fixation to identify all CB1R, we differentiated cells expressing CB1R predominantly at the plasma membrane (PM-_UbC_SSF-CB1) from those at endo-membranes (EM-_UbC_SSF-CB1). This allowed us to separate both populations post-imaging by cell sorting and to separate their unique cAMP responses. In the PM-_UbC_SSF-CB1 population, we observed an increase in cAMP levels after WIN55,212 treatment and a decrease of cAMP in the EM-_UbC_SSF-CB1 population (Figure 2B,C). These findings underscore that the sub-cellular localization of CB1R significantly dictates its downstream signaling, particularly its impact on cAMP levels. Our findings suggest that CB1R positioned at the plasma membrane primarily couples with Gs proteins, leading to increased cAMP levels, whereas localization in endo-membranes results in Gi/o coupling and decreased cAMP levels (Table 1).

**Table 1:** Location bias of CB1R leads to differences in cAMP signaling.

| promoter | N-ter tag | predominant localization | cAMP signaling |
| --- | --- | --- | --- |
| CMV | WT | PM | increase cAMP |
| UbC | WT | EM | decrease cAMP |
|  | SSF | PM | increase cAMP |
|  |  | EM | decrease cAMP |

### Location bias: EM-CB1R prefers Gαi/o, PM-CB1R prefers Gαs

Next, we investigated whether these effects were directly mediated by G-protein coupling or through alternative mechanistic pathways. Heterotrimeric G proteins and adenylated cyclase have been observed not only associated to the plasma membrane but also to intracellular compartments such as endosomes or the Golgi, supporting the concept of endo-membrane based G-protein signaling(10, 11, 31, 32). Recently, it has been demonstrated that the β1-adrenergic receptor and the opioid receptor mu can both localize at the Golgi and activate Gαs and Gαi respectively (10, 11). To assess the role of G proteins in CB1R-mediated cAMP signaling, we generated siRNA-based knockdowns of specific G proteins in our cell model (Figure S2.2). We first performed a Gαs knockdown and co-expressed the EPAC sensor with _UbC_SSF-CB1, the receptor construct which expresses at low levels but is primarily at the cell surface due to the non-native signal sequence (SS). After treatment with the CB1R agonist, WIN55,212, we observed a delayed and significantly smaller increase in cAMP levels compared to non-pretreated cells (Table 1, Figure 2E,F), indicating that PM-CB1R predominantly activates Gαs for cAMP signaling although a limited contribution of Gαi cannot be excluded.

Next, we generated a Gαi knockdown and co-expressed the EPAC sensor with _UbC_CB1. Consistent with our observations, knockdown of Gαi blocked all effects of WIN55,212 treatment in the endo-membrane localized _UbC_CB1 cells (Figure 2D,F). To further confirm this finding, we treated cells with Pertussis toxin for 24h, a treatment known to induce ADP-ribosylation and subsequent degradation of Gαi/o. Interestingly, in Pertussis toxin-treated cells expressing _UbC_CB1, we observed an increase in cAMP levels after WIN55,212 treatment (Figure 2D,F). We believe that this increase is driven by the small fraction of CB1R at the plasma membrane that still couples with Gαs in _UbC_CB1 expressing cells. The results from the Gαs knockdown and Pertussis toxin experiments support the idea that PM-CB1R predominantly activates Gαs for cAMP signaling. Correspondingly, knockdown of Gαi/o blocked CB1R effects on cAMP in cells in which CB1R was largely retained at endomembranes. Our findings indicate that the sub-cellular localization of CB1R plays a crucial role in determining its downstream signaling through specific G protein coupling. In our model system in which we can control CB1 localization with the SSF tag without measurably affecting expression levels, PM-CB1R largely activates Gαs, leading to an increase in cAMP levels, while EM-CB1R mostly activates Gαi, resulting in decreased cAMP levels.

### Minimally invasive labeling of CB1R by a non-canonical amino acid as an alternative method to distinguish EM-CB1R to PM-CB1R

We found that a standard tag used for monitoring GPCR expression and trafficking, SSF, affected CB1R localization and signaling. However, the simple solution to this problem -- removing SSF tag -- would make it highly challenging to monitoring CB1R trafficking in living cells. We therefore aimed to develop a novel approach for monitoring CB1R expression and trafficking by using the smallest possible tagging technique while maintaining the ability to distinguish between cells with CB1R predominantly at the plasma membrane or in intracellular compartments. We sought to achieve this goal without interfering with cannabinoid receptor signaling or trafficking. To accomplish this, we used genetic code expansion to incorporate a *trans*-cyclooct-2-en-L-lysine (TCO*A) for catalyst-free ultrafast labeling of the receptor, a technique previously employed for other membrane proteins.(33) We utilized an orthogonal tRNA/tRNA synthetase pair (tRNA/RS) from *Methanosarcina mazei* to introduce TCO*A lysine into the first extracellular loops of CB1R (Figure 3A). We used a CB1-GFP construct to screen several positions for the most receptor expression and efficient TCO*A incorporation, and the most successful site was found to be by replacing the phenylalanine at position 180 in CB1R (_CMV_CB1-F180)(Figure 3B and 4A). We also added an SSF tag at the N-terminus of _CMV_CB1-F180 to assess its functionality, specifically looking at the kinetics of internalization after treatment with WIN55,212 (10µM). For labeling CB1 before agonist treatment, we utilized the non-cell-permeable dye methyl tetrazine ATTO 647 (ATTO 647 MeTet). The functionality of _CMV_CB1-F180 was assessed by examining its internalization kinetics compared to _CMV_SSFCB1 (wild-type CB1R with an SSF tag) after treatment with WIN55,212. Both _CMV_SSFCB1 and _CMV_SSFCB1-F180 showed similar levels of receptor internalization (approximately 75% and 70%, respectively) during 3 hours of treatment (Figure 3C,D,K). This suggest that the F180 tag minimally perturbed CB1R functionality. To further confirm the functional integrity of _CMV_CB1-F180, we activated the receptor by uncaging caged-2AG, which has previously been shown to transiently increase intracellular calcium levels ([Ca^2+^]_i_) (34). Uncaging showed a response at the same order of magnitude as through wild-type CB1R expression (Figure 3E). Notably, the response to 2AG was completely abolished by the CB1R inverse agonist rimonabant, validating the specificity of _CMV_CB1-F180 (Figure 3E,F,G).

**Figure 3.**
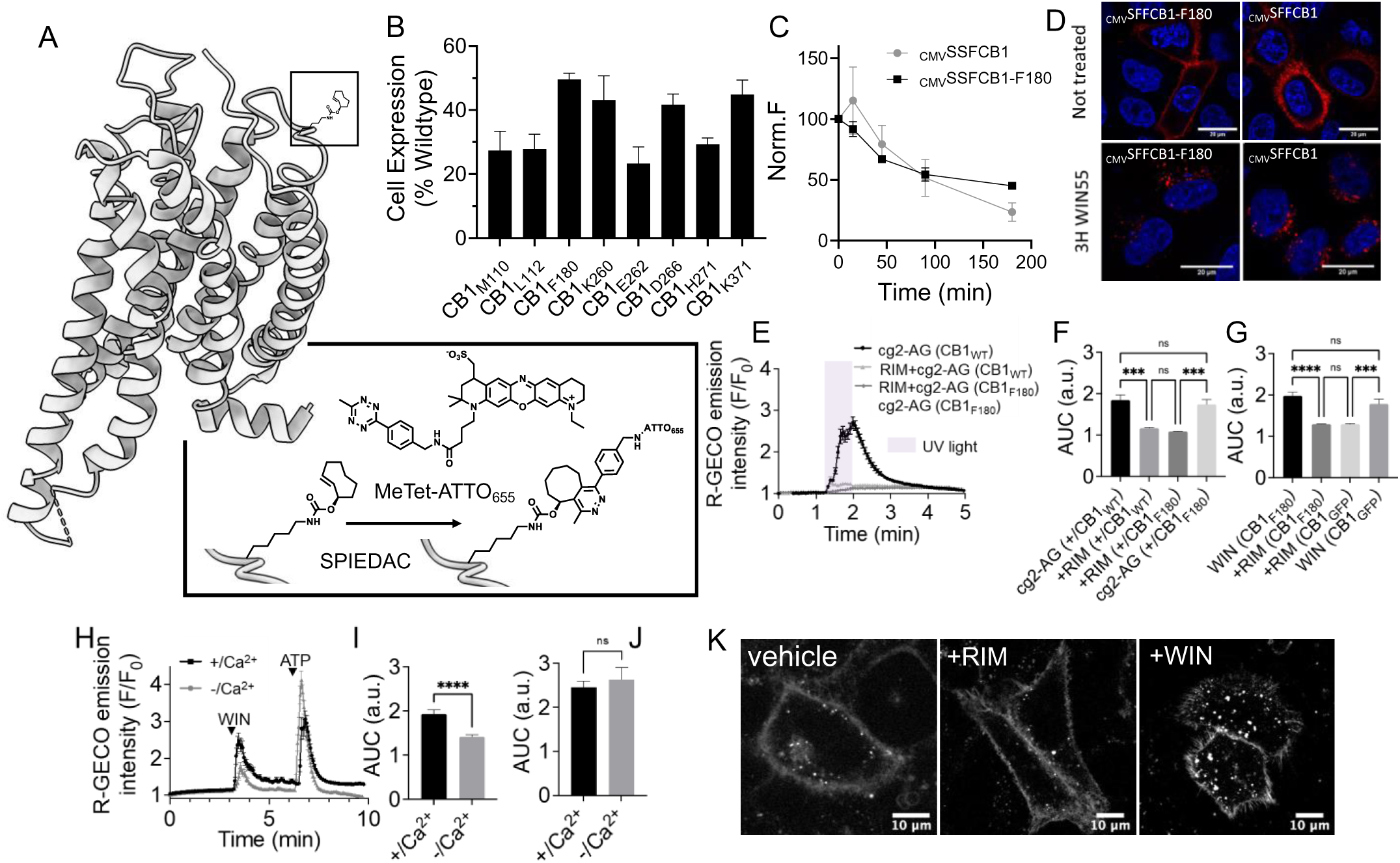
Introducing a clickable TCO*A in CB1R without interfering with signaling or trafficking provides an alternative method to distinguish EM-CB1R to PM-CB1R. **A.** Structural model of CB1R having the unnatural amino acid TCO*A incorporated before and after SPIEDAC reaction with the methyl tetrazine dye Me-Tet-ATTO_655_. **B.** Single cell analysis of GFP fluorescence following transient transfection of CB1-GFP that contained TCO*A at the position indicated. Data represent the mean +/− SD of biological triplicate. The fluorescence was detected with a Beckman Coulter Cytoflex S. **C.** Time course of WIN55,212-2 (10 µM) stimulated receptor internalization as assayed by loss of cell surface immunoreactivity and measured by flow cytometry comparing un-modified _CMV_SSFCB1 with the _CMV_SSFCB1-F180. (10 µM WIN55, 0,15,45,90 and 180 min after stimulation). **D.** Confocal micrographs showing live HeLa Kyoto cells transfected with _CMV_SSFCB1 or _CMV_SSFCB1-F180. Cells were immuno-labeled with a functionalized Flag antibody and treated for 3 h with WIN55. **E.** Fluctuations of [Ca^2+^]_i_ measured by R-GECO in CB1-WT or CB1-F180-transfected HeLa Kyoto subjected to treatment with the CB1 agonist cg2-AG (10 µM) and illuminated by 375nm UV light, pretreated or not with the antagonist rimonabant. **F,G.** Comparison of area under the curve fluorescence of R-GECO in CB1-WT or CB1-F180-transfected HeLa Kyoto after uncaging cg2-AG (10 µM) under different experimental conditions. **H.** Fluctuations of [Ca^2+^]_i_ in CB1R-transfected HeLa Kyoto upon treatment with WIN55,212-2 (10 µM) and with ATP (50 µM), pre-incubated or not in nominally Ca^2+^-free media. **I.** Comparison of area under the curve fluorescence of R-GECO in CB1R-transfected cells after treatment with WIN55,212-2 (10 µM). **J.** Comparison of area under the curve fluorescence of R-GECO in CB1R-transfected cells after treatment with ATP (50µM). **K.** Confocal micrographs showing HeLa Kyoto cells transfected with CB1-F180, tagged with MeTet-ATTO655 and treated with the CB1R antagonist rimonabant (10µM) or the agonist WIN55,212-2 (10µM).

Our alternative method for tagging CB1R model by incorporating the TCO*A label at position 180 in CB1R enabled efficient and catalyst-free ultrafast labeling of the receptor exclusively at the plasma membrane, allowing discrimination from intracellular compartment localization. Importantly, functional assessment demonstrated minimal perturbation of CB1R by the F180 tag, as evidenced by normal internalization kinetics and intact calcium signaling responses to 2AG.

### Effects of CB1 expression and activation on calcium levels and receptor internalization

To control for potential contributions of intracellular calcium levels to cannabinoid signaling, we employed a classical approach and expressed a GFP-fused version of CB1R with an unmodified N-terminus under the CMV promoter along with the calcium sensor R-GECO to monitor changes in intracellular calcium concentrations ([Ca^2+^]_i_) upon CB1R activation. WIN55,212-2 triggered a transient increase in [Ca^2+^]_i_ (F/F_0_=2.545±0.128, n=80, Figure S3A,B). This response was completely abolished by pre-treating cells with the inverse agonist rimonabant (F/F_0_=1.267±0.003, n=170, Figure S3A,B). In control experiments, ATP addition induced a major calcium transient (F/F_0_=3.339±0.119, n=89), likely through Gq/11-coupled P2Y receptors (Figure S3A,B)(35). Intracellular calcium stores are known to be involved in the CB1R-mediated increase in cytoplasmic calcium levels in different cellular models(13, 36, 37). This was confirmed by a short incubation with thapsigargin, a non-competitive inhibitor of the sarco/endoplasmic reticulum Ca^2+^ ATPase (SERCA) (F/F_0_=1.761±0.131, n=80), or Xestospongin C (an IP_3_ receptor antagonist) respectively, which reduced the calcium response to WIN55,212-2 (Figure S3C,D). This result suggests that CB1R-mediated calcium increase partially relies on the release of calcium from internal stores mediated by IP_3_ and IP_3_ receptors. As expected, both thapsigargin and xestospongin C also dramatically decreased the cytosolic increase in calcium after addition of ATP (Figure S3C,E). To explore if extracellular calcium intake via GIRK channels is a primary pathway after CB1R activation, we incubated cells in Ca^2+^-depleted media supplemented with EGTA for 5 min. The reduction in calcium response after WIN55,212 addition in calcium-depleted media suggested that both extracellular intake and release from intracellular stores contributed to the cytosolic increase in calcium levels (Figure 3H,I). Importantly, the extracellular depletion of calcium had no impact on the cytosolic increase in calcium after addition of ATP (Figure 3H,J).

We also observed that prolonged activation of CB1R triggered receptor internalization and co-localization with Rab5 in early endosomes. Desensitization of the receptor upon internalization was indicated by the inability of a second dose of the agonist to provide another transient calcium response (F/F_0_=1.107±0.008, n=80, Figure S3K,L). In contrast, inactivation of CB1R with the reverse agonist rimonabant led to the accumulation of the receptor at the plasma membrane (Figure S3F,I,J). In summary, CB1R activation induced calcium release from intracellular stores, likely through IP_3_ receptors, and extracellular calcium influx. The desensitization of CB1R upon prolonged activation highlights the receptor’s critical regulatory role in maintaining cellular responses.

### Cannabinoid receptor constructs regulate adenylate cyclase activity and trigger distinct signaling pathways

Next, we aimed to investigate how the newly developed _CMV_CB1-F180 construct regulates adenylate cyclase (AC) activity and to compare it’s signaling to the unmodified _CMV_CB1 receptor. To assess adenylate cyclase (AC) activity, we co-transfected _CMV_CB1-F180 or _CMV_CB1 with the EPAC-based sensor and measured cAMP levels after stimulation with WIN55,212 (10 µM) followed by forskolin (50µM) (Figure 4A,B,C). Both _CMV_CB1-F180 and _CMV_CB1 expressing cells showed a comparable moderate increase in intracellular cAMP levels after WIN55,212 stimulation. However, the difference in cAMP levels between _CMV_CB1-F180 and _CMV_CB1 was more pronounced after forskolin stimulation, with _CMV_CB1-F180 expressing cells displaying moderately higher cAMP levels than wild type cells. These results indicate that _CMV_CB1-F180-mediated signaling is comparable to the unmodified receptor and that _CMV_CB1-F180 predominantly couples to Gαs proteins (Figure 4B,C). Furthermore, the _CMV_CB1-F180 construct showed approximately 50% expression level relative to unmodified _CMV_CB1 (Figure 3B). This led us to hypothesize that two populations exist in _CMV_CB1-F180 transfected cells: one with lower expression level and CB1R predominantly at endo-membranes and another with higher expression level and CB1R present in both plasma and endo-membranes. By post-translationally labeling _CMV_CB1-F180 before experiments with the non-cell-permeant dye methyl tetrazine ATTO 647 (ATTO 647 MeTet) and tagging of all CB1R receptors post fixation with a CB1R-specific antibody, we distinguished between cells with CB1R predominantly at the plasma and endo-membranes (PM-_CMV_CB1-F180) and exclusively in endo-membranes (EM-_CMV_CB1-F180)(Figure S4.2). After CB1R activation, PM-_CMV_CB1-F180 cells showed an increase, while EM-_CMV_CB1-F180 cells displayed a decrease in cAMP levels, indicative of Gαs and Gαi/o coupling, respectively (Figure 4D,E). These data provide a parallel line of evidence that cells in which CB1 predominantly localizes to internal membranes primarily couples to Gαi/o while cells in which CB1 is largely at the cell surface signal through Gαs.

**Figure 4.**
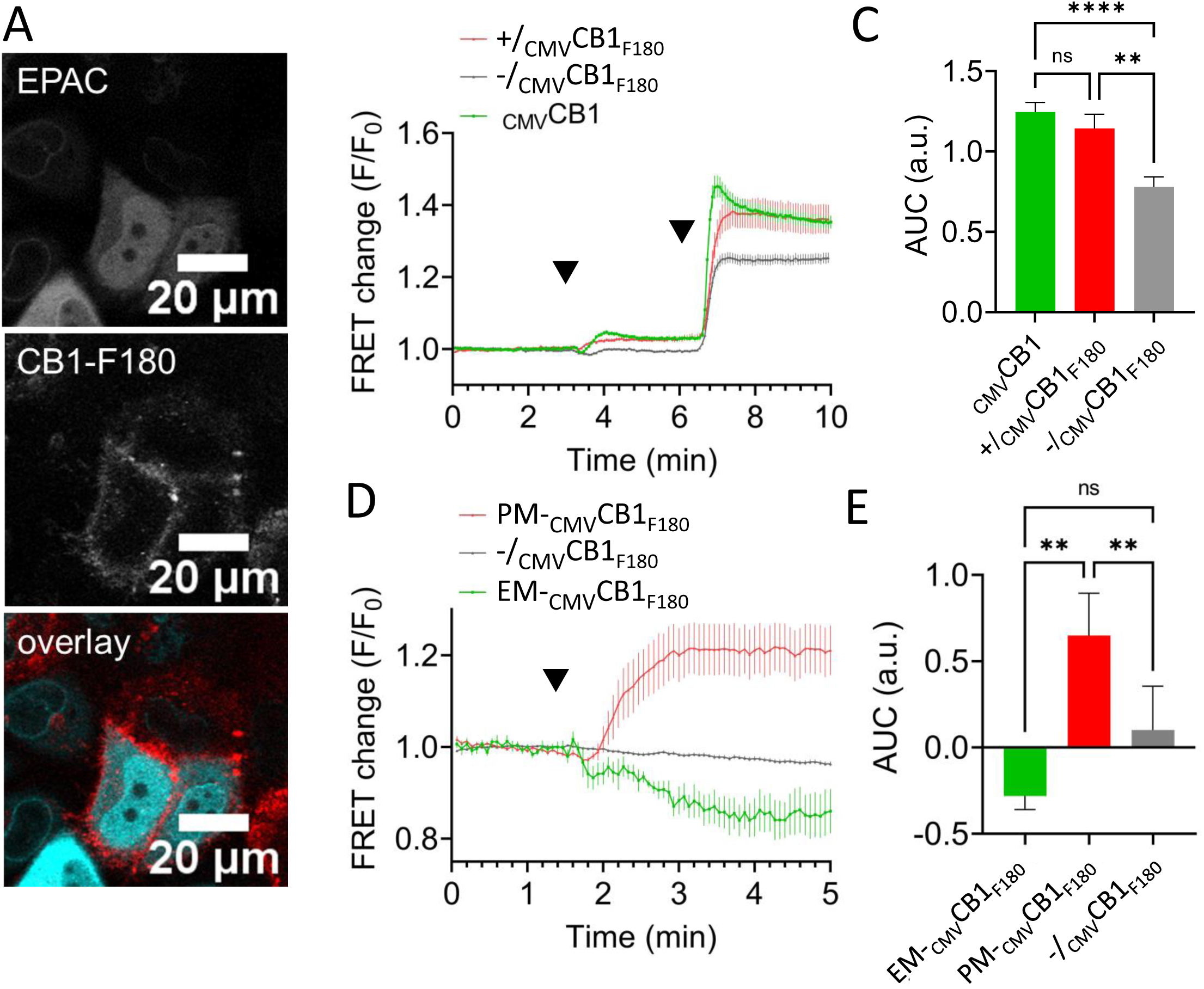
_CMV_CB1-F180 and CB2-S29 constructs regulate AC activity and demonstrate that distinct signaling patterns depend on CB1 localization. **A.** Confocal micrographs showing _CMV_CB1-F180 and EPAC FRET sensor co-transfected HeLa Kyoto cells after receptor labelling with Me-Tet-ATTO655 (1 µM) for 20 min. **B.** Average of 45, 178 and 76 cell traces showing FRET changes of the EPAC-based FRET sensor in _CMV_CB1-F180-transfected HeLa (+/), (−/) and wt _CMV_CB1 respectively, after treatment with WIN55,212-2 (10 µM) followed by forskolin (FSK, 50 µM). **C.** Bar graphs showing FRET changes of the EPAC-based FRET sensor after forskolin stimulation (50 µM) in −/_CMV_CB1-F180 (WT) versus +/_CMV_CB1-F180 p < 0.005, _CMV_CB1 versus +/_CMV_CB1-F180 non-significant or _CMV_CB1 versus −/_CMV_CB1-F180 p < 0.001expressing cells. **D.** Average of 14, 6 and 82 cell traces showing FRET changes of the EPAC-based FRET sensor in _CMV_CB1-F180-expressing cells tagged with Me-Tet-ATTO655 (+PM-_CMV_CB1-F180), _CMV_CB1-F180-expressing cells without me-Tet-ATTO655 (EM-_CMV_CB1-F180) and non-expressing cells (−/_CMV_CB1-F180) respectively, after treatment with WIN55,212-2 (10 µM). **E.** Bar graphs showing area under the curve of the EPAC-based FRET sensor after WIN55,212-2 stimulation (10 µM) in −/_CMV_CB1-F180 non-expressing cells, PM-_CMV_CB1-F180 or EM-_CMV_CB1-F180 expressing cells. EM-_CMV_CB1-F180 versus PM-_CMV_CB1-F180 p < 0.005, PM-_CMV_CB1-F180 versus −/_CMV_CB1-F180 p < 0.005, EM-_CMV_CB1-F180 versus −/_CMV_CB1-F180 was non-significant.

As a control, we also incorporated TCO*A lysine in the N-terminus of the CB2 receptor (CB2R) at serine 29, termed CB2-S29, to explore its G-protein coupling and signaling characteristics (Figure S4.1A). CB2R is a class A GPCR sharing 44% homology with CB1R and can be activated by WIN55,212 as a full agonist. It predominantly couples to Gαi/o proteins and unlike CB1R, has shown little promiscuity for other G proteins. We observed no changes in cAMP levels after WIN55,212 stimulation in CB2-S29 expressing cells compared to non-transfected cells (Figure S4.1B,C). However, after forskolin treatment, CB2-S29 cells showed lower cAMP levels, suggesting predominantly Gαi/o coupling, consistent with CB2R being a Gαi/o-coupled receptor. When treated with the CB2-inverse agonist AM630, CB2-S29 showed significantly higher level of [cAMP]_I_ after forskolin stimulation compared to non-transfected cells (Figure S4.1B,D) suggesting a strong Gαi/o interaction (Figure S4.1B,C). Our study provides valuable insights into the signaling characteristics of _CMV_CB1-F180 and CB2-S29 receptors. _CMV_CB1-F180 retains its ability to signal predominantly via Gαs coupling, while CB2-S29 exhibits Gαi/o coupling. Additionally, our results support the idea that CB1R can signal from endo-membranes and modulate its G-protein coupling based on its subcellular localization.

## Discussion

Recent developments in the area of GPCR molecular pharmacology have shown that GPCRs can signal from endomembranes—such as endosomes and Golgi—as well as the plasma membrane. We set out to examine how CB1R signaling is dependent on subcellular location and how spatial bias could contribute to cannabinoid activity. Our data strongly indicate that CB1R from internal membranes predominantly couples to Gαi/o while CB1R at the cell surface prefers to couple to Gαs, suggesting an encoding of spatial bias in cannabinoid receptor function (Figure 5). A critical step in our finding was developing methods to control, and deconvolve, CB1R subcellular localization. We have demonstrated that CB1R trafficking and location can be highly dependent on its expression level and modification of the CB1R N-terminal tail with a non-native epitope tag (SSF). This allows for control of CB1R localization. Importantly, we provide an alternative solution to the commonly used SSF for tracking cannabinoid receptor trafficking by developing methods for single site incorporation of *trans*-cyclooctene lysine (TCO*A) into the extracellular loops of CB1R or CB2R.

**Figure 5.**
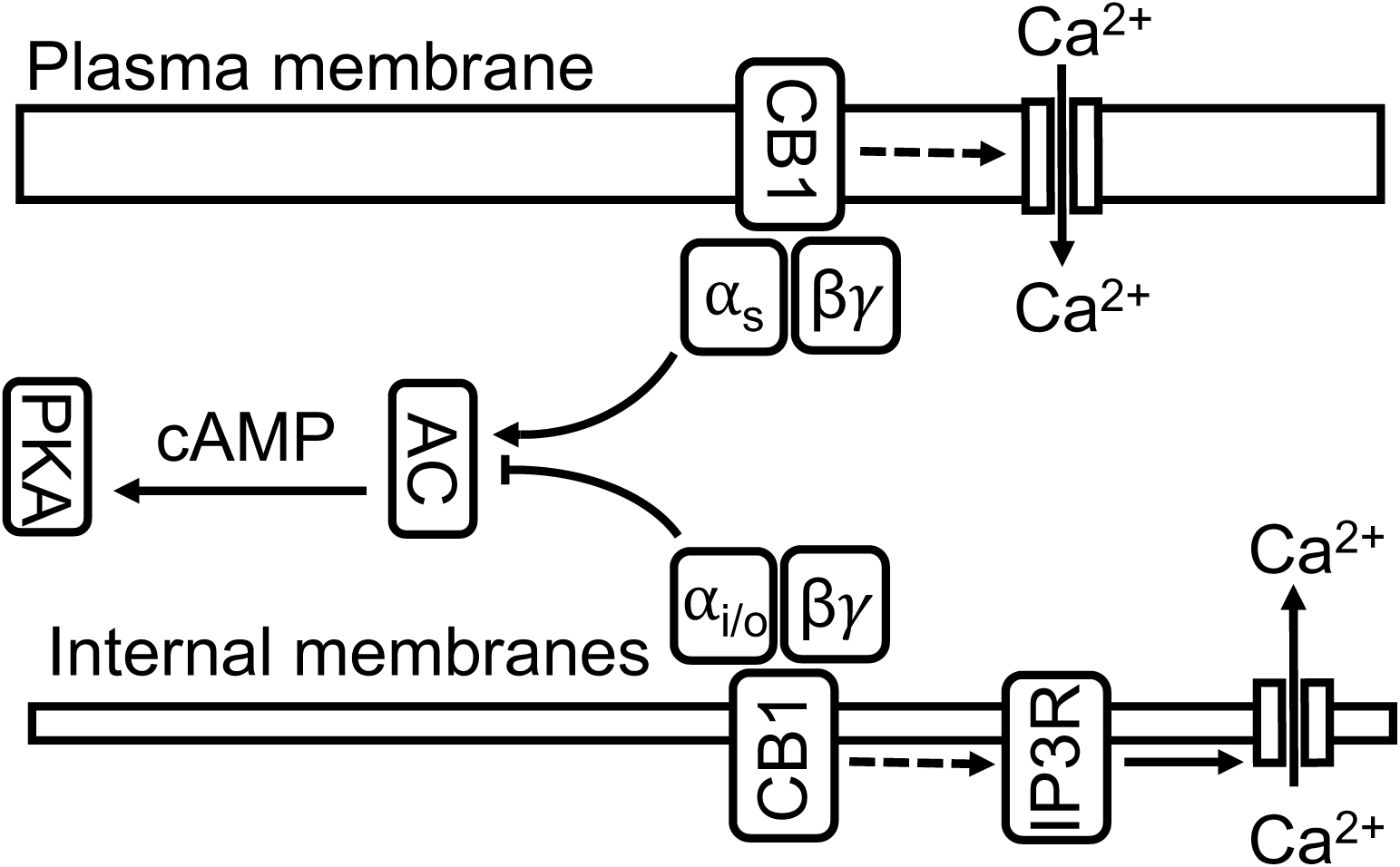
The scheme depicts the contradictory effect of activated CB1 on cAMP levels depending on receptor location.

### Subcellular Localization of CB1R

The subcellular localization of CB1R has been a point of controversy. It is mostly believed that functional CB1R receptors are present at the plasma membrane and ligand binding there is fully responsible for its signaling activity (38, 39). Some studies have shown that the internal pool is a result of CB1R internalization from the plasma membrane and trafficking to internal membranes (40, 41). It is believed that CB1R is cycling back to the plasma membrane after inverse-agonist treatment (42). This view has been challenged by other studies demonstrating that internalized CB1R is mostly degraded in the endolysosomal pathway and that the internal pool does not contribute to the plasma membrane CB1R population (43). We found in this study that CB1R, at relative low expression levels, resides predominantly in internal membranes with very low plasma membrane expression. While we did not characterize the mode of trafficking, our findings suggest that the internal pool of CB1R in proximity to the Golgi was never trafficked to the plasma membrane. This aligns with the studies describing internal CB1R as an independently operating receptor pool. In our model, expression levels are the main driver of basal sub-cellular localization suggesting that the formation of the CB1R intracellular pool is potentially a result of retention by an adaptor protein preventing CB1R to enter trafficking vesicles and to reach the plasma membrane, and such a mechanism has been described for several other GPCRs (44, 45).

### Location Bias in CB1R Signaling

We demonstrate, using different expression and tagging systems, that CB1R signaling is highly influenced by its subcellular localization: plasma membrane CB1R can predominantly couple to Gαs and promotes cAMP accumulation while intracellular CB1R shows dominant coupling to Gαi and inhibits cAMP production. Our findings indicate that agonist stimulation leads to a differential response in cAMP levels based on CB1R localization: an increase via Gαs coupling in cells with surface-expressed CB1R, and a decrease through Gαi signaling in cells where CB1R is primarily located in endo-membranes. Measurements of intracellular cAMP levels in combination with a predominant location of CB1 at internal membranes through a weak promoter suggest that CB1R can signal from internal membranes and is subject to location bias. This result has major implications on the pharmacology of CB1 stimulation. Specifically, the lipophilicity of a ligand, or its synthesis location in the case of endocannabinoids, will influence its preferential binding to either the plasma membrane or endomembrane population of the receptor. Our observations raise the question, why does CB1R predominantly couple to Gαs at the cell surface and Gαi at endomembranes? Recent studies have suggested that the membrane itself plays a key role in GPCR coupling to G proteins. Specifically, anionic phospholipids at the plasma membrane like PI(4,5)P_2_ have been shown to control coupling of the GPCR B2AR to G proteins through charge-based suppression of the receptor interaction with Gαi (46). Building on this finding, recent studies show that this lipid-based regulation can give rise to location bias in B2AR signaling as the Golgi is not enriched in PI(4,5)P_2_ (47). Thus, we speculate that one of the underlying mechanisms for our observation of location bias in CB1R signaling is differential phospholipids enrichment at endomembranes compared to the cell surface. Finally, overexpressed CB1R has been previously shown to couple not only to Gαi/o but also to Gαs (29). It was argued that the presence of a “receptor reserve” amplifies GPCR signaling and makes low affinity binding of Gαs appear as the main response(48). We cannot exclude that the CB1R coupling to Gαs is in part a product of this phenomena. It will be important in the future to use methods to restrict CB1R activation or visualization in order to clearly identify CB1R signaling, effector activation and ligand accessibility(32, 49).

### Role of CB1R N-terminus in its Subcellular Localization

The CB1 N-terminal tail is uniquely long within the rhodopsin receptor family and has been suggested to play a role in its expression, trafficking and signaling(50). However, the function of the N-terminus is still poorly understood. As was already shown previously(28, 41), we demonstrated that modifying the N-terminal portion even by only adding an SSF tag dramatically affects CB1R’s sub-cellular localization. This turned out to be an effective tool in allowing for deconvolving the role of subcellular localization in CB1R signaling.

### CB1R Signaling via Calcium

The signaling picture is not complete without accounting for CB1R-induced calcium transients. Our results show that there is a component of WIN-induced calcium peaks that is driven by calcium influx which could be triggered by Gβγ at the plasma membrane. However, there is a second component that is sensitive to thapsigargin and therefore originates likely from intracellular calcium stores. Whether this is induced by Gαq(13) or through calcium-induced calcium release remains unclear.

### Minimally Preturbative Methods for Monitoring Cannabinoid Receptor Trafficking

We have demonstrated that tags commonly used for monitoring GPCR expression and trafficking, like SSF, perturb the subcellular localization of CB1R. To provide a solution to this issue, we have developed an alternative method for live cell tagging and tracking of CB receptors. We used genetic code expansion in combination with strain-promoted inverse electron-demand Diels-Alder chemistry (SPIEDAC) which provides a minimal alteration and changes the labeling from the N-terminus to the first extracellular loop. This proved to be an efficient method for tagging the receptor at the plasma membrane while having little impact on CB1R signaling and trafficking, and we identified sites for efficient incorporation into both CB1R and CB2R. Previously, other groups have employed genetic code expansion to study GPCRs and other membrane receptors in a variety of settings (33, 51). We believe our SPIEDAC incorporation site will allow for high flexibility in the choice of dye and will hence provide a valuable tool for future studies of CB1R in intact cells.

In summary, we demonstrated that expression level and N-terminal modification of CB1R can lead to disruption in the receptor location and function. We identified a modulatory signal transduction of CB1R dependent on the receptor’s cellular location, indicative of location bias. Consequently, the synthesis of agonists and antagonists with cell-permeable or impermeable properties, designed to target intracellular organelles or to bind receptors exclusively at the cellular surface, respectively, may be instrumental in leveraging the functional selectivity of the CB1R.

## METHODS

### RESOURCE AVAILABILITY

#### Lead Contact

Further information and requests for resources and reagents should be directed to and will be fulfilled by the Lead Contact, Aurélien Laguerre

#### Materials Availability

This study did not generate new unique reagents.

#### Data and Code Availability

This study did not generate any unique datasets or code.

### EXPERIMENTAL MODEL AND SUBJECT DETAILS

The HeLa Kyoto cell line (RRID:CVCL_1922, female) was kindly provided by R. Pepperkok (European Molecular Biology Laboratory, Germany). HeLa Kyoto (passage 15-35) were grown in 4.5g/L glucose DMEM (Life Technologies, 41965-039) supplied with 10 % fetal bovine serum (Life Technologies, 10270098).

### METHOD DETAILS

#### General

All chemicals were obtained from commercial sources (Acros, Sigma-Aldrich, Tocris, TCI, Cayman, Alfa Aesar, Atto-tec or Merck) and were used without further purification unless otherwise specified. Rimonabant (5-(4-chlorophenyl)-1-(2,4-dichlorophenyl)-4-methyl-N-(piperidin-1-yl)-1H-pyrazole-3-carboxamide), AM630 ((6-iodo-2-methyl-1-(2-morpholinoethyl)-1H-indol-3-yl)(4-methoxyphenyl) methanone, Forskolin (5-(acetyloxy)-3-ethenyldodecahydro-6,10,10b-trihydroxy-3,4a,7,7,10a-pentamethyl-(3R,4aR,5S,6S,6aS,10S,10aR,10bS)-1H-naphtho[2,1-b]pyran-1-one), Xestospongin C ([1R-(1R,4aR,11R,12aS,13S,16aS,23R,24aS)]-eicosahydro-5H,17H-1,23:11,13-diethano-2H,14H-[1,11]dioxacycloeicosino[2,3-b:12,13- b1]dipyridine) and WIN55, 212-2 ([(11R)-2-methyl-11-(morpholin-4-ylmethyl)-9-oxa1-azatricyclo[6.3.1.04,12]dodeca-2,4(12),5,7-tetraen-3-yl]-naphthalen-1-ylmethanone) from Cayman Chemical were dissolved in dimethylsulfoxide (DMSO) to a stock concentration of 10 mM. Thapsigargin ((3S,3aS,4R,6R,7S,8R)-6-acetoxy-4-(butyryloxy)-3,3a-dihydroxy-3,6,9-trimethyl-8-(((Z)-2-methylbut-2-enoyl)oxy)-2-oxo-2,3,3a,4,5,6,6a,7,8,9b-decahydro-1H-cyclopenta[e]azulen-7-yl octanoate) from Sigma was dissolved in DMSO to a stock concentration of 5 mM. ATP (adenosine 5’-triphosphate disodium salt hydrate) from TCI was freshly dissolved in DMSO to a concentration of 10 mM. Atto488 Me-Tetrazine from Atto-Tec was dissolved in DMSO to a stock concentration of 1 mM. cg2-AG was synthesized, purified and chemically characterized following the methods previously reported in the literature (for details, see Laguerre A, Hauke S, Qiu J, Kelly MJ, Schultz C. Photorelease of 2-Arachidonoylglycerol in Live Cells. *J Am Chem Soc.* **2019**;141(42):16544–16547. doi:10.1021/jacs.9b05978). All chemicals were administrated to cells with a DMSO concentration lower or equal to 0.1 %.

#### Amplex intact cell assay

Cells were seeded in a 6 well plates. After 24 h, CB1-APEX constructs were transfected according to the Lipofectamine 2000 (Life Technologies, 11668030) manufacturer protocol. After overnight incubation the transfection medium was replaced with fresh full growth medium. 24 h post transfection cells were lifted and resuspended in PBS. AUR (Amplex UltraRed, Thermo, A36006) was added to cells from a 10 mM stock to a 2 µM final concentration and incubated at RT for 5 min followed by 10 min at 4°C (from there all steps were performed at 4°C). Cells were then incubated in PBS supplemented with 2% BSA and 50 µM H_2_O_2_ for 1 min. The reaction was quenched with 1 mM sodium azide, the cells were spined down and washed with PBS + 2% BSA. Immediately after cells were imaged on a FACS. The AUR fluorescent reaction product was detected with a Beckman Coulter Cytoflex S (excitation 633, emission 670/30).

#### Immunostaining of HeLa cells

After incubation with transfection mix or microscopy, cells were fixed with 4% paraformaldehyde for 15 min, washed twice in PBS, permeabilized in 0.1% Triton X-100 for 2 min and immunostained with primary antibodies overnight. The cells were then washed four time in PBS and incubated with a secondary antibody for 1 h. Cells were then washed four times with PBS and imaged on a dual scanner confocal microscope Olympus Fluoview 1200, using a 63x (oil) objectives.

#### Genetic code expansion and SPIEDAC tagging

Cells were seeded in eight-well Lab-Tek microscope dishes for 24 h (to reach 60-70 % confluence) before transfection. After 24 h, 200 ng of hMbPylRS-4xU6M15 (Addgene, #105830) and 200 ng of the respective amber construct were premixed in 20 µL of DMEM. 0.3 µL of Lipofectamine 2000 (Life Technologies, 11668030) in 20 µL of DMEM was then added to the DNA premix and incubated for 20 min at RT before being added to the wells. Shortly after the transfection mixture was added to cells, 100 µM of the ncAA TCOA*K was added from a 100 mM stock solution in 0.1 M NaOH. After overnight incubation, the transfection medium was replaced with fresh full growth medium. 30 min before imaging, cells were washed two times with DMEM (without FBS) and incubated for 20 min with 1 µM of Me-Tet Atto488 from a 1 mM stock solution in DMSO. After 20 min cells were washed with imaging medium (Invitrogen, A14291DJ) four times before imaging.

#### Calcium imaging experiments

Cells were seeded in eight-well Lab-Tek microscope dishes for 24 h (to reach 60-70 % confluence) before transfection. For imaging of CB1-GFP transfected cells, 100 ng of CB1-GFP and 100 ng of R-GECO (Addgene #32444) were mixed with 0.2 µL of lipofectamine 2000 transfection reagent. For imaging of cells transfected with CB1-F180 or CB2-S29, the experimental protocol described in the genetic code expansion section was followed with an addition of 200 ng of R-GECO. For all of the above mixes, DNAs and lipofectamine were separately premixed in 20 µL of DMEM then mixed together and incubated for 20 min before being added to each well of the eight well Lab-Tek containing 200 µL of DMEM 4.5g/L glucose supplemented with 10% FBS. Cells were imaged at 37°C in imaging buffer. Imaging was performed on a dual scanner confocal microscope Olympus Fluoview 1200, using a 63x (oil) objective. The R-GECO sensor was imaged using a 559 nm laser (at a laser power of 1.0%) and a 643/50 emission filter. Fluctuations of [Ca^2+^]_i_ were monitored through excitation at 559 nm and emission above 600 nm (F/F_0_) on the confocal microscope.

#### Trafficking experiments

Cells were seeded in eight-well Lab-Tek microscope dishes for 24 h before transfection. 100 ng of CB1-GFP and 100 ng of Rab5-BFP (Addgene #49147) were mixed with 0.2 µL of lipofectamine 2000 following transfection method previously described then added to the wells. 48 h after the transfection, cells were incubated with 10 µM of WIN55,212-22 or 10 µM of rimonabant for 3 h. Cells were imaged at 37°C in imaging buffer. Imaging was performed on a dual scanner confocal microscope Olympus Fluoview 1200, using a 63x (oil) objectives.

#### EPAC-based sensor imaging experiments

Cells were seeded in eight-well Lab-Tek microscope dishes for 24 h (to reach 60-70 % confluence) before transfection. 100 ng of the EPAC sensor (Addgene #61622) were mixed with 0.2 µL of lipofectamine 2000 following the transfection method previously described. After overnight incubation the transfection medium was replaced with fresh full growth medium. 24 h after the first transfection, CB1R (100 ng) or CB2R (100 ng) were mixed with 0.2 µL of lipofectamine 2000 and added to cells. For CB1-F180 or CB2-S29, the experimental protocol described in the genetic code expansion section was used after the cells were first transfected with the EPAC sensor. Cells were imaged at 37°C in imaging buffer. Imaging was performed on a dual scanner confocal microscope Olympus Fluoview 1200, using a 63x (oil) objective. The EPAC FRET sensor was imaged using a 440 nm laser (at a laser power of 1.0%) and the signal was collected in the CFP/YFP emission channels.

To sort between cells expressing CB1R at the plasma membrane from cells expressing CB1R in endomembrane, the exact field of view during the live cell experiment was saved and the cells removed from the microscope stage and fixed in 4% PFA for 15 min at RT. The cells were then immunostained following the protocol described above. After immunostaining the cells were reset on the microscope stage and using the saved coordinate, re-imaged using an identical field of view and sorted by hand using ImageJ.

#### siRNA knockdown assay

siRNA (20µM) was mixed with dharmafect according to the manufacturer protocol recommendation for HeLa Kyoto. Cells were seeded at 10% confluence with transfection mix. After overnight incubation, the transfection medium was replaced with fresh full growth medium. For microscopy, the EPAC-based sensor imaging protocol was performed 48 h after cell seeding. For blotting, cells were lysed 80 h post seeding in RIPA buffer (Thermo, 89901) containing cOmplete EDTA-free protease inhibitor cocktail (Roche). Lysate were centrifuged at 15000 rpm for 15 min. Total protein was quantified with a BCA assay (Thermo, 23225), normalized, denatured in sample buffer by boiling for 5 min at 95 °C and resolved in a 4-12% gradient gel Bis-Tris gel (Thermo, NP0321BOX). The proteins were then transfer to a PVDF membrane using Trans-Blot Turbo Transfer system (Bio-Rad), blocked in 3% milk PBST and incubated overnight at 4°C with primary antibodies. Blots were washed with PBST 4×5min and incubated with secondary HRP conjugated antibodies for one hour and washed 4×5min. Developing was done with SuperSignal West Femto Maximum Sensitivity Substrate (Thermo, 34095) and imaged on a ChemiDoc (BioRad).

#### Images analysis

All Images were analyzed on the FIJI software using the pipeline summarized in Figure S1B. Primarily, multi-channel images were separated into single channels and converted to 8-bit for calcium imaging or 32-bit for EPAC experiments. The time course experiment was duplicated and stacked using the Z project function (RFP channel for calcium imaging and CFP channel for EPAC imaging). Using the paintbrush tool set at 0, cells were manually delimited to achieve robust single cell segmentation. A mask of regions of interest was generated using the combination of the threshold and analyze particles tools (as depicted in Figure S1B). This ROI mask was then superimposed to the time course experiment and the multi-measure function was applied to it. From this stack, we extracted mean single cell values from the time course experiment. Those values were then exported to an excel files for further analysis.

#### Statistical analysis

All statistical comparisons were performed using One way ANOVA by Prism or Excel. Statistical details of each experiment can be found in the figures and figure legends. For all experiments, the number of cells and error bars (SEM) can be found in the results section and the respective figure legends. All imaging experiments were performed at least in biological triplicates, n indicating the total number of cells.

### KEY RESOURCES TABLE

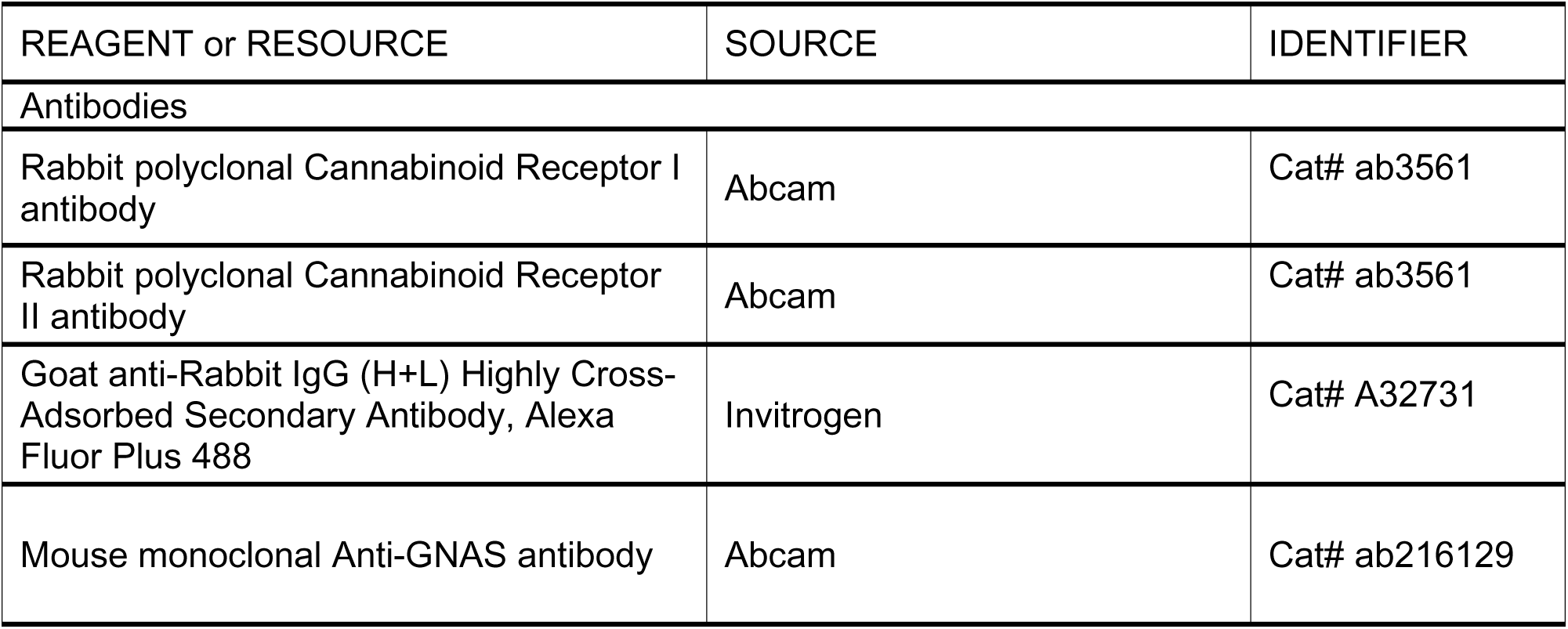
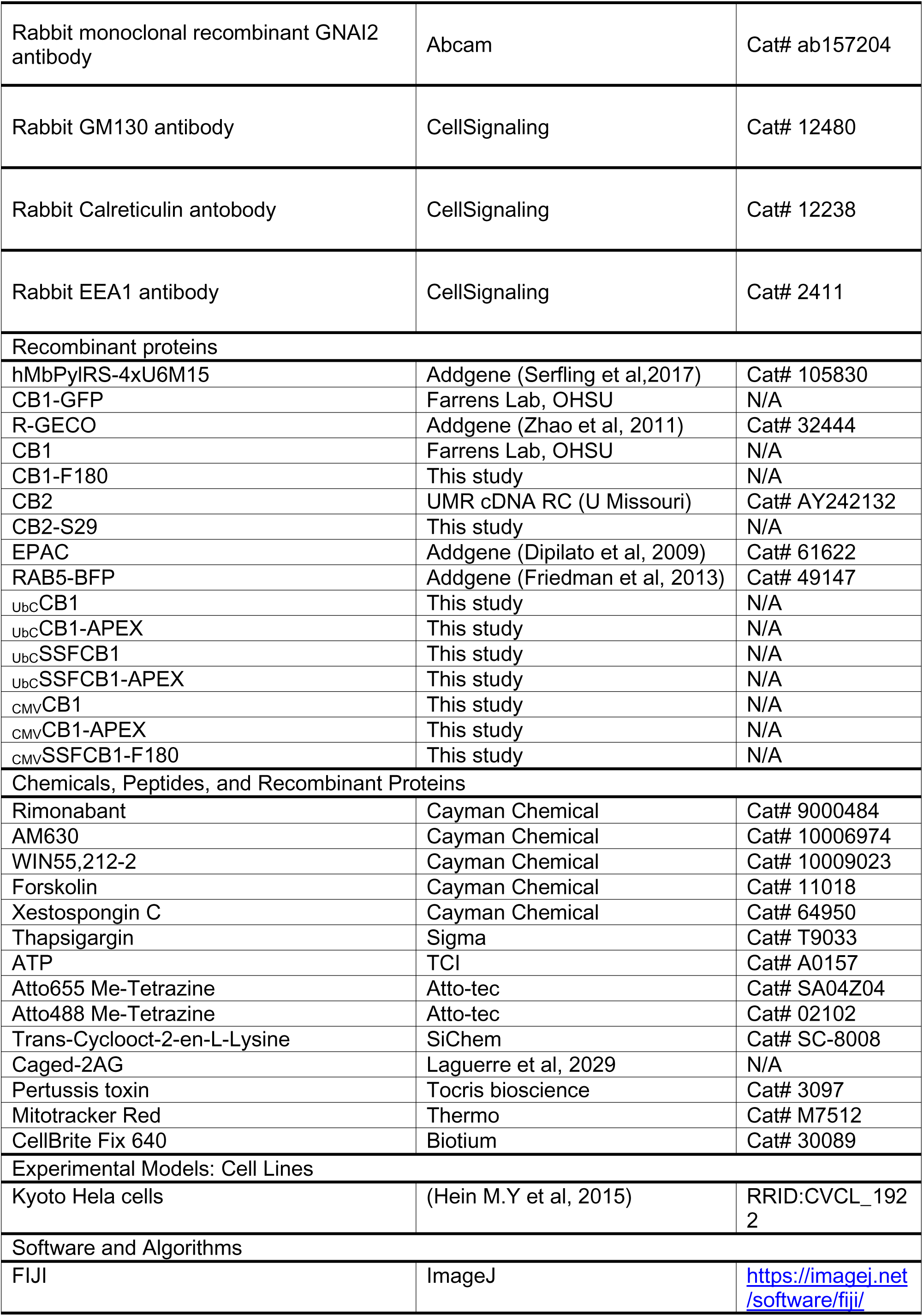
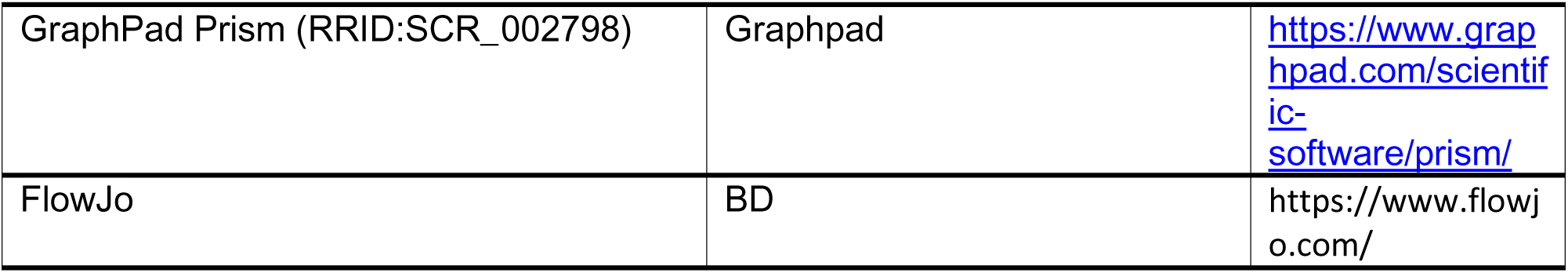

## ACKNOWLEDGEMENTS

The authors thank the Farrens group from OHSU for providing wild type CB1R. A.T., C.S. and A.L. acknowledge financial support from OHSU. B.L was supported by NIH grant GM137835, C.S. by GM127631. A.L. was supported by K99 GM141316. C.S. is a recipient of a Mercator Fellowship from the DFG, connected to Transregio 186.

## AUTHOR CONTRIBUTIONS

A.T. designed the study, performed cloning and imaging experiments, analyzed data, interpreted the results and co-wrote the manuscript. C.S. and B.L. interpreted the results and co-wrote the manuscript. A.L. designed the study, performed imaging experiments, analyzed data, interpreted the results and co-wrote the manuscript.

---

**Figure S1.1.**
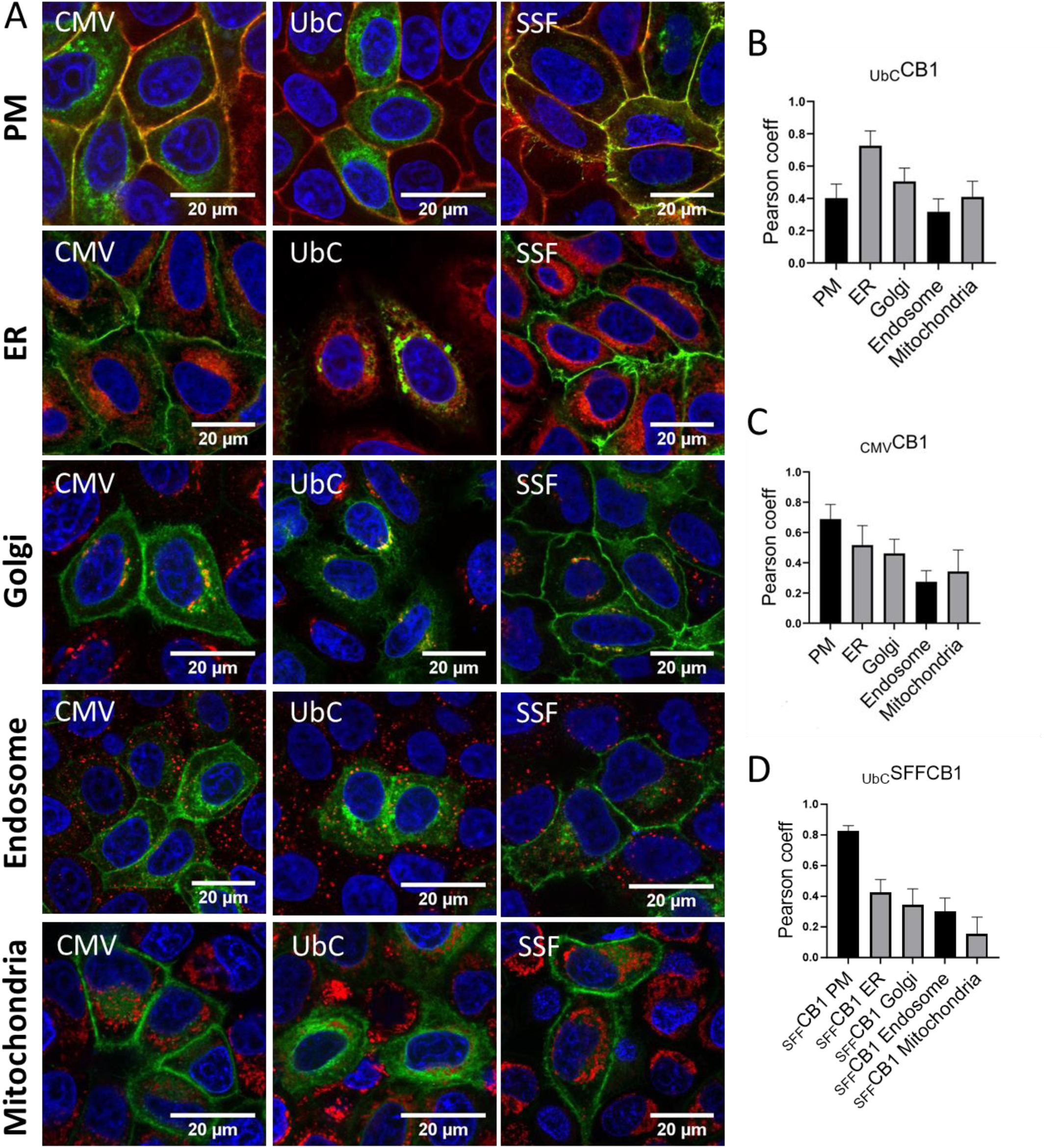
**A.** Confocal micrographs showing HeLa Kyoto cells transfected with _UbC_CB1R, _CMV_CB1R or _UbC_SSFCB1R. Cells were fixed, permeabilized and c-immuno-stained with a CB1R antibody and the PM, ER, Golgi, endosome and mitochondria organelle markers cellBrite, Calreticulin, GM130, EEA1 and MitoTracker, respectively. Co-localization measurement in HeLa cells using Pearson coefficient of immunolabeled _UbC_CB1R (**B**), of immunolabeled _CMV_CB1R (**C**) or immunolabeled _UbC_SSFCB1R (**D**) with respective organelle markers.

**Figure S1.2.**
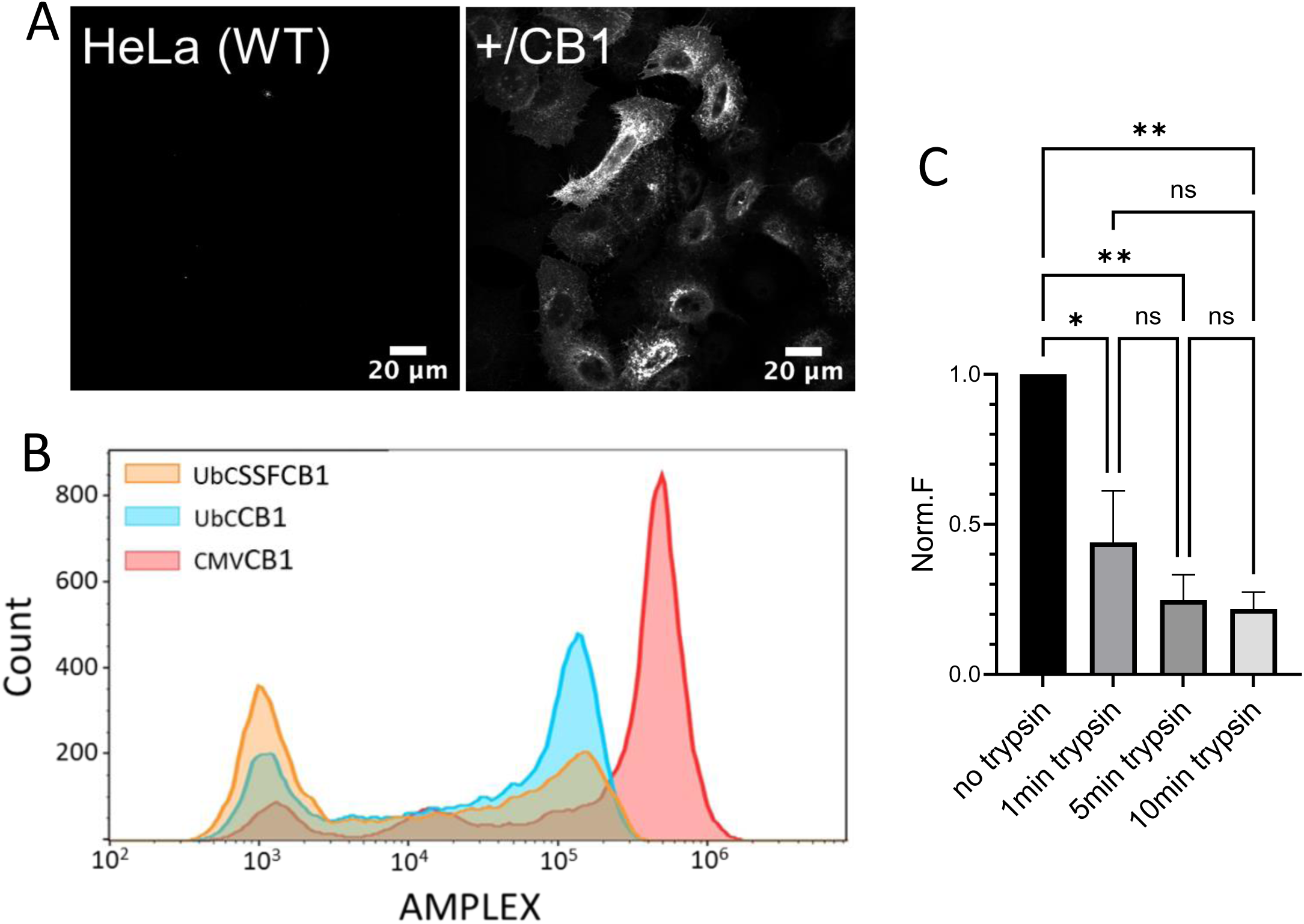
**A.** Immuno-fluorescence experiment comparing CB1R expression between untransfected (WT) or _CMV_CB1R (+/CB1) HeLa cells. **B.** Single cell analysis of APEX2/AUR intact assay following transient transfection of either _CMV_CB1R-APEX2, _UbC_CB1R-APEX2 or _UbC_SSFCB1R-APEX2. The fluorescence reaction product of the APEX2 reaction with AUR was detected with a Beckman Coulter Cytoflex S. **C.** Time course bar graph of _UbC_SSFCB1 as assayed by loss of cell surface immunoreactivity and measured by flow cytometry comparing non-trypsinized _UbC_SSFCB1R-expressing cells with trypsinized _UbC_SSFCB1R cells. (0,1,5,10 min time points).

**Figure S2.1.**
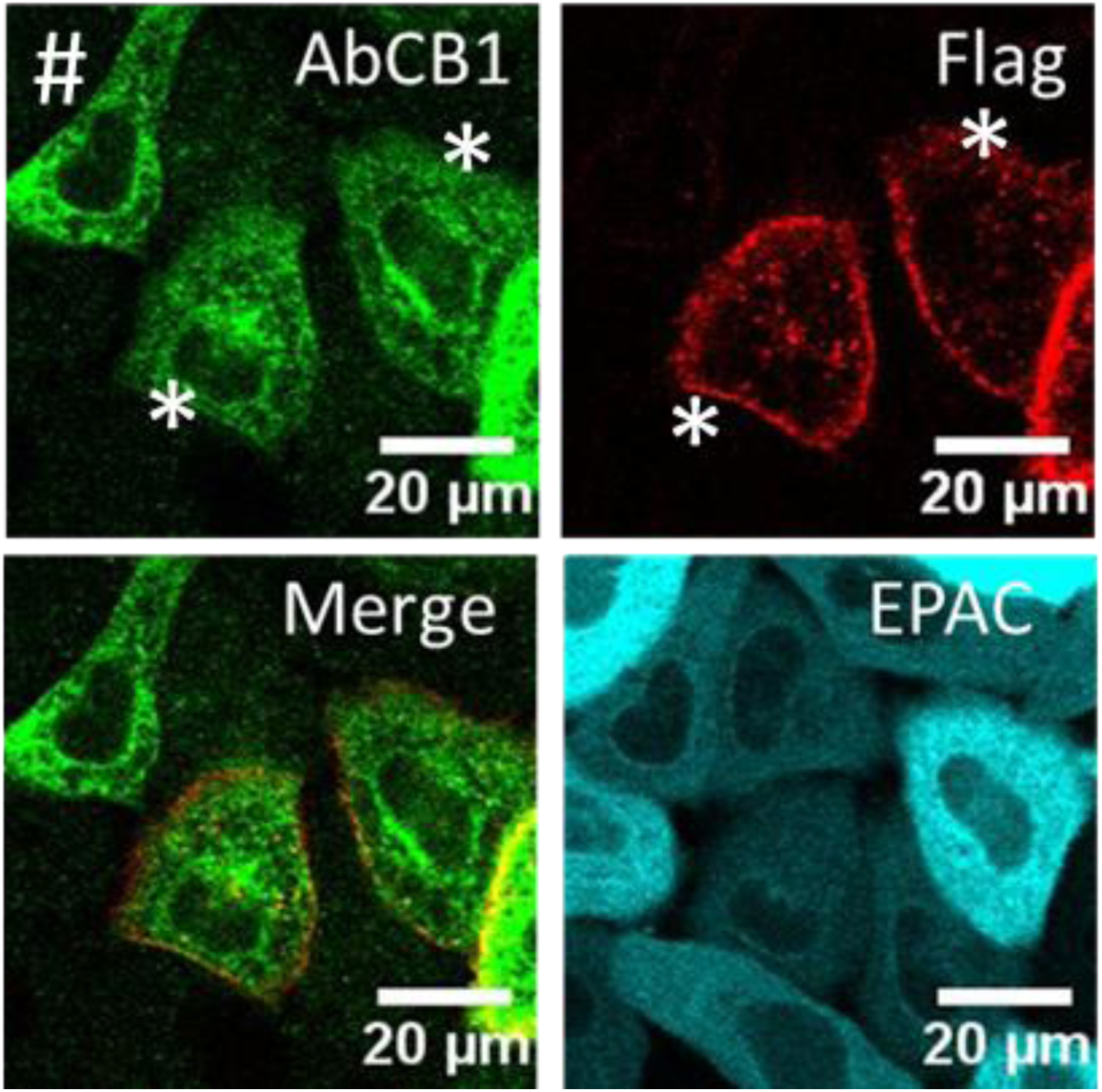
Confocal micrographs showing HeLa Kyoto cells transfected with _UbC_SSFCB1R. Receptors were labelled with a functionalized 647Alexa-M1-FLAG antibody for 20 min in live cells (top right), then fixed, permeabilized and immuno-stained with a CB1R antibody (top left). Note that cells marked with a (#) are labeled exclusively by the CB1 antibody post permeabilization while cells marked with a (*) are labeled with both CB1R and Flag antibody

**Figure S2.2.**
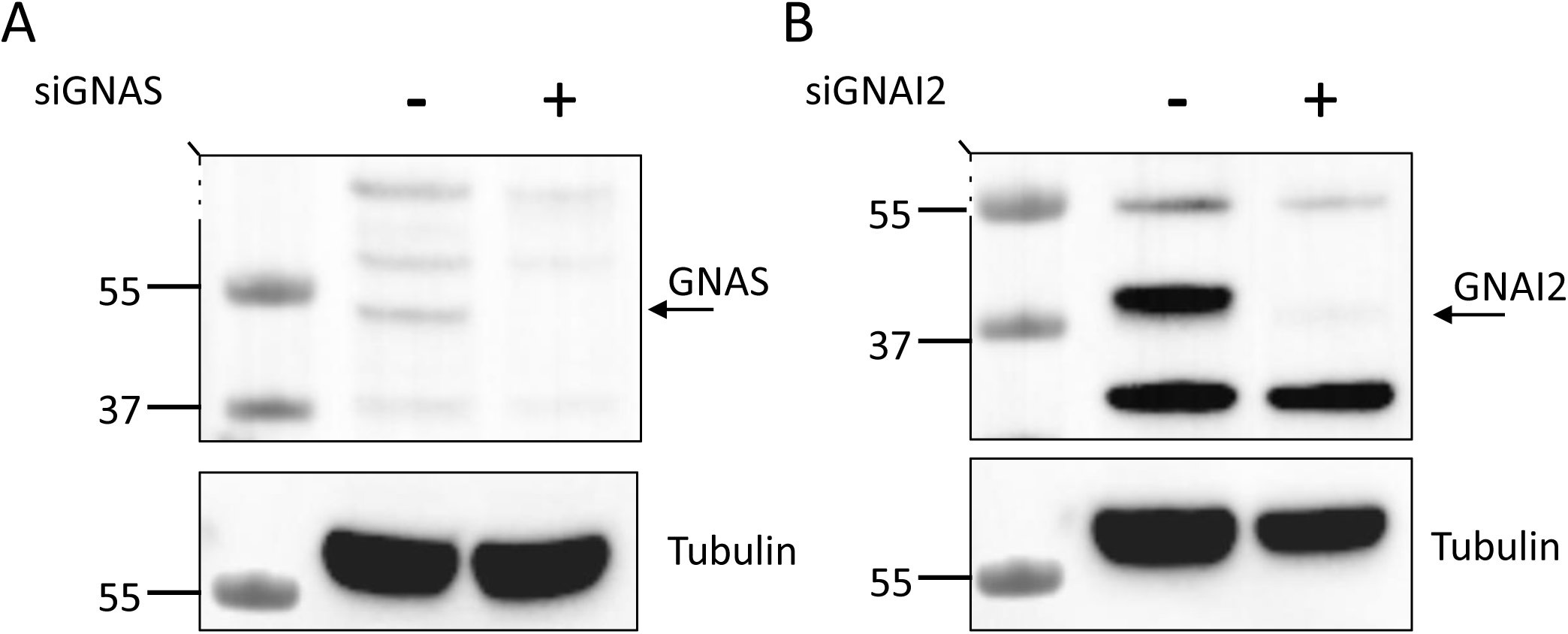
Immunoblot analysis of HeLa Kyoto cells upon **A.** Gαs (*GNAS*) knockdown and **B.** Gαi (*GNAI2*) knockdown. Arrows indicate the corresponding molecular weight of *GNAS* and *GNAI2*. (N=2). Dash lines indicate slicing.

**Figure S3.**
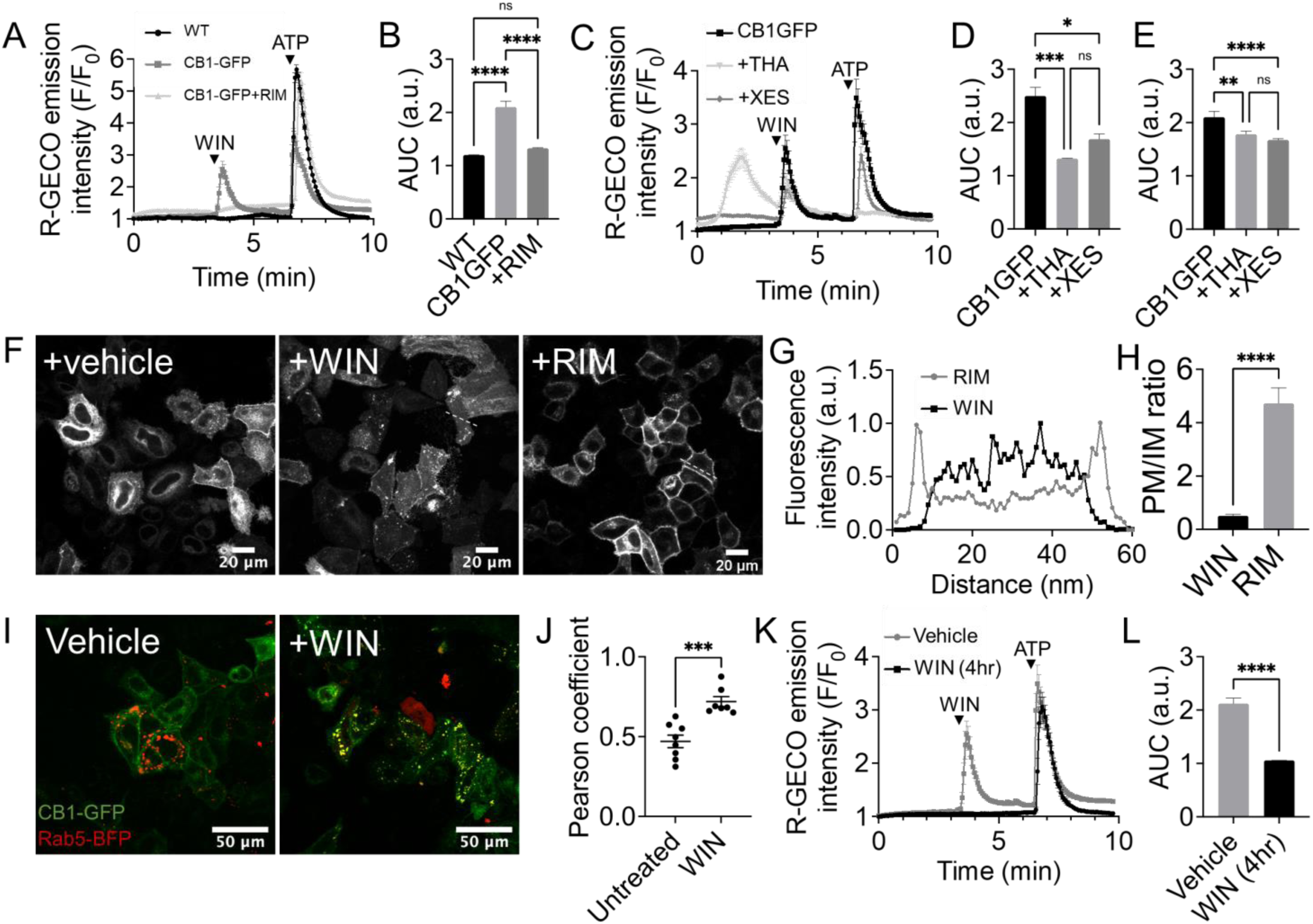
CB1R activation leads to a transient increase in calcium levels via influx of extracellular calcium and intracellular release. **A.** Fluctuations of [Ca^2+^]_i_ in CB1-GFP transfected HeLa Kyoto upon treatment with WIN55,212-2 (10µM) and with ATP (50µM) pre-treated or not with the antagonist rimonabant. **B.** Comparison of area under the curve fluorescence of R-GECO in CB1-GFP transfected HeLa Kyoto after treatment with WIN55,212-2 (10µM) under different experimental conditions. **C.** Fluctuations of [Ca^2+^]_i_ in CB1-GFP transfected HeLa Kyoto upon treatment with WIN55,212-2 (10µM) and with ATP (50µM) pre-treated or not with Thapsigargin (10µM) or Xestospongin C(10µM). **D.E.** Comparison of area under the curve fluorescence of R-GECO in CB1-GFP transfected HeLa Kyoto after treatment with WIN55,212-2 (10µM) and with ATP (50µM) pre-treated or not with Thapsigargin (10µM) or Xestospongin C (10µM). **F.** Confocal micrographs (20x) showing CB1-GFP transfected cells after 3 h treatment with 10 µM of WIN55,212-2 or with 10 µM of rimonabant. **G.H.** To quantify the ratio of CB1-GFP at the plasma membrane to the endomembrane after different treatments a line was drawn across each cell and the fluorescence across the line was measured. **I.** Confocal micrographs showing the relative subcellular locations of CB1-GFP and Rab5-BFP with vehicle or with WIN55,212-2 (10 µM) for 3 hr. **J.** Colocalization measurement in HeLa cells using Pearson coefficient of CB1-GFP and Rab5-BFP. **K.** Fluctuations of [Ca^2+^]_i_ in CB1-GFP transfected HeLa Kyoto upon treatment with WIN55,212-2 (10µM) and with ATP (50µM) pre-treated or not for 4hours with WIN55,212-2 (10µM). **L.** Comparison of area under the curve fluorescence of R-GECO.

**Figure S4.1.**
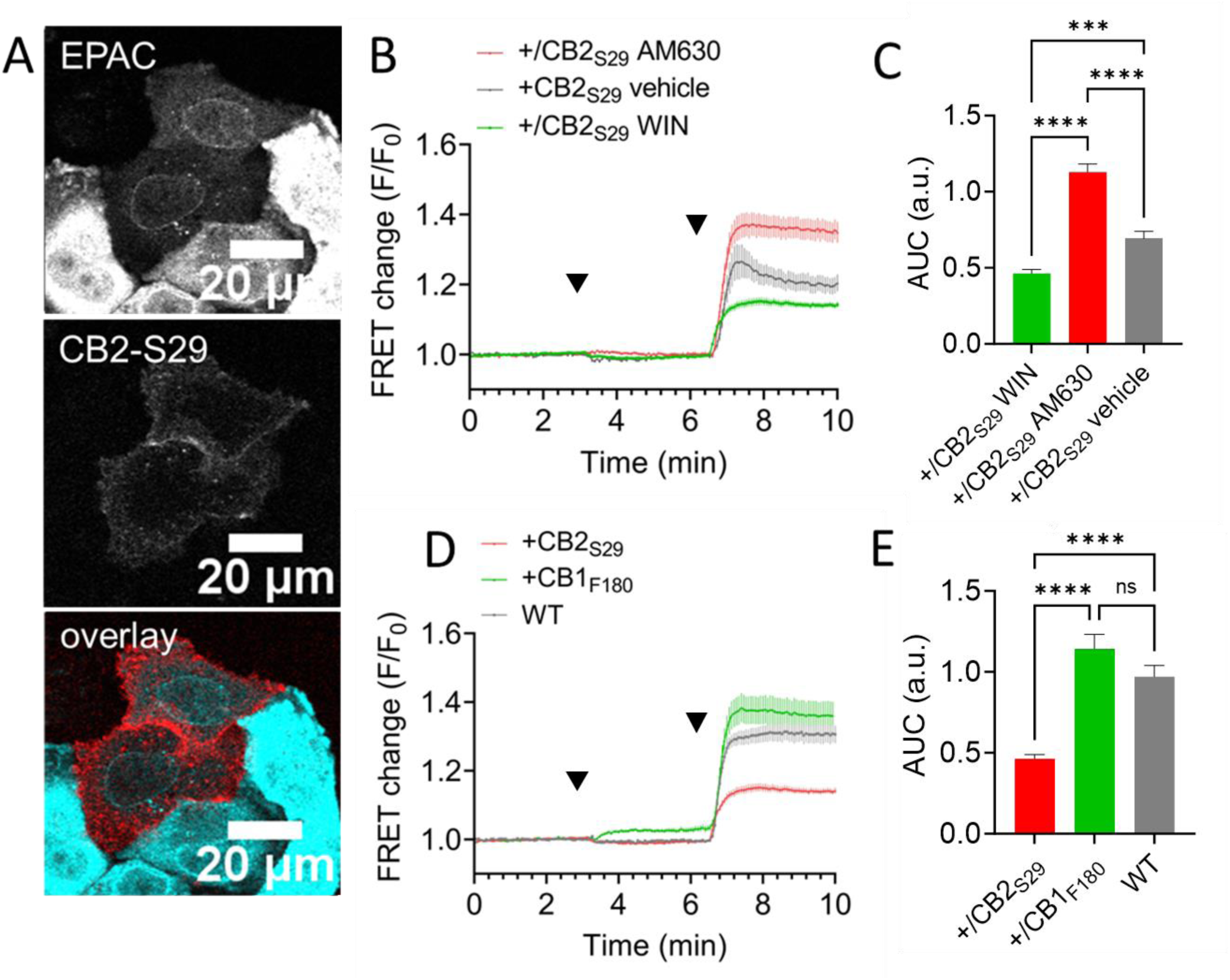
CB1-F180 and CB2-S29 constructs regulate AC activity and demonstrate distinct signaling pathways. **A.** Confocal micrographs showing HeLa Kyoto cells co-transfected with CB2-S29 and the EPAC-based FRET sensor. Receptors were labelled with Me-Tet-ATTO655 (1 µM) for 20 min. **B.** Average of 21, 16 and 27 cell traces showing FRET changes of the EPAC-based FRET sensor in CB2-S29-transfected HeLa, after treatment with the agonist WIN55,212-2 (10 µM) or the inverse agonist AM630 (1 µM), followed by forskolin (FSK, 50 µM). **C.** Bar graphs showing FRET changes of the EPAC-based FRET sensor after forskolin stimulation (50 µM) in CB2-S29 expressing cells tagged with Me-Tet-ATTO655 (+/CB2R). WIN - vehicle p < 0.005, WIN vs AM630 p < 0.001, AM630 vs vehicle p < 0.001. **D.** Average of 27, 45 and 70 cell traces showing FRET changes of the EPAC-based FRET sensor in CB2-S29 and CB1-F180-transfected HeLa, after treatment with the agonist WIN55,212-2 (10 µM), followed by forskolin (FSK, 50 µM). **E.** Bar graphs showing FRET changes of the EPAC-based FRET sensor after WIN55,212-2 and forskolin stimulation ((50 µM) in wild type (WT) versus CB2-S29 expressing cells tagged with Me-Tet-ATTO655 (+/CB2-S29) and CB1-F180 expressing cells (+/CB1-F180). +/CB2R versus +/CB1R p <0.0001, +/CB1R versus WT p-value non-significant.

**FigureS4.2.**
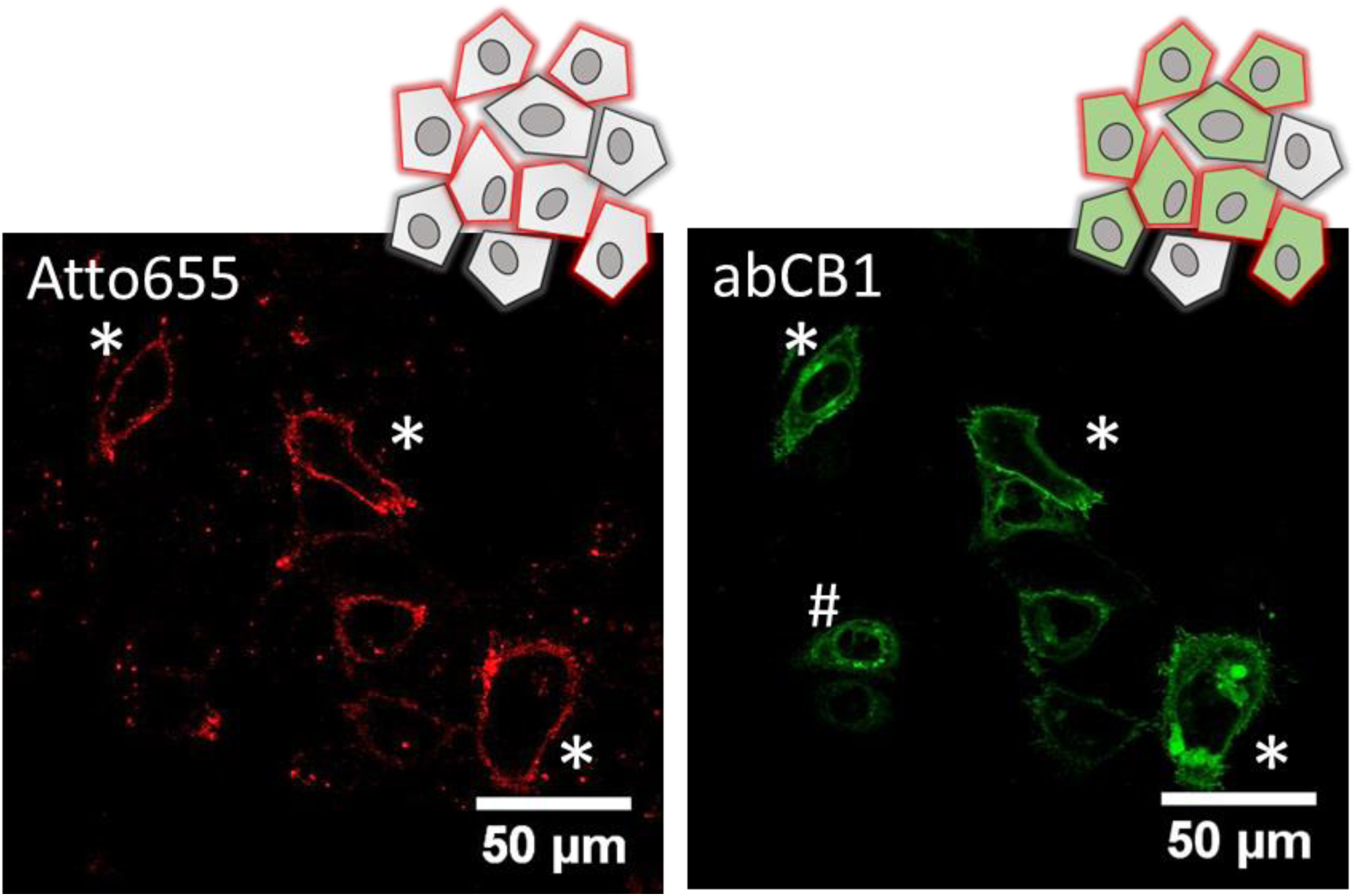
Additional evidence from CB1-F180 post-translational labeling and change in [cAMP] levels shows that CB1R localization in endo-membrane affects G-protein coupling. Confocal micrographs showing HeLa Kyoto cells transfected with CB1-F180. Receptors were labelled with Me-Tet-ATTO655 (1 µM) for 20 min (left panel) then fixed and immuno-stained with a CB1R antibody. Note that cells marked with a (#) are labeled exclusively by the antibody while cells marked with a (*) are labeled with both antibody and Me-Tet-ATTO655. This experiment is representative of three biological repeats.

